# Sperm storage reduces sperm and embryo quality in animals

**DOI:** 10.1101/2025.04.10.647966

**Authors:** Krish Sanghvi, Rebecca Dean, Shinichi Nakagawa, Klaus Reinhardt, Irem Sepil, Regina Vega-Trejo

**Author notes:** Shared first authors- equal contribution (Krish Sanghvi, Rebecca Dean).

## Abstract

In many animals, sperm are stored for extended periods either in the reproductive tracts of males before ejaculation, or of females after copulation. Sperm storage reduces the risk of sperm limitation in both sexes and avoids the costs of female re-mating. However, sperm storage can lead to post-meiotic sperm senescence, i.e. within-sperm-age-dependent deterioration, potentially impacting conceived offspring and lowering male and female fitness. Yet, the extent and magnitude of such deterioration and the variables modulating it during sperm storage are not well understood. Using a meta-analysis across humans (115 studies) and non-human animals (56 studies from 30 species), we investigate how in-vivo sperm storage affects sperm quality, fertilisation success, and offspring quality. In humans, sperm storage leads to greater sperm oxidative stress and DNA damage, and reduces sperm viability and motility. In other animals, sperm performance and embryo quality decline. We identify the duration of sperm storage, the design used for sampling individuals, and the sex of the individual storing sperm as potentially important moderators of the effects of sperm storage. These findings have key biomedical implications, including optimising the timing of ejaculation and fertilisation in fertility clinics or captive breeding programs. Overall, our results reveal the mechanisms that cause post-meiotic sperm senescence, the fitness consequences of sperm storage, and provide evolutionary insights into sex-specific adaptations that potentially mitigate the detrimental effects of sperm storage.

## Introduction

Across the animal kingdom, males store mature spermatozoa for extended durations between spermatogenesis and ejaculation, while internally fertilising females do so between mating and fertilisation (Orr and Brennan, 2015; Pizzari et al, 2008; Reinhardt, 2007). For instance, males of some bat (Crichton, 2000; Sato et al, 2023) and frog (Hettyey et al, 2012) species store mature sperm over many months of hibernation; while queen Hymenoptera (Den Boer, et al, 2009) and females of some reptiles (Neubaum and Wolfner, 1998) store sperm for many years. In males, sexual rest enables the storage and accumulation of sperm inside the reproductive tract, potentially minimising the risk of sperm limitation under polygyny or sperm competition (Orr and Brennan, 2015; Sanghvi et al, 2024a, 2025). In females, sperm storage helps reduce re-mating for obtaining additional sperm when mating is harmful, provides the opportunity for effective oviposition site selection, facilitates sperm competition and cryptic female choice, and prevents sperm depletion when males are rarely encountered (Birkhead and Moller, 1993; Holt and Lloyd, 2010; Jimenez-Franco et al, 2020; Orr and Brennan, 2015; Orr and Zuk, 2012). However, sperm storage in males and females can lead to the deterioration of sperm quality and even impact the fitness of the conceived offspring, due to post-meiotic sperm senescence (reviewed in Du et al, 2024; Pizzari et al, 2008; Reinhardt, 2007; Tarin et al, 2000).

There are several physiological reasons why individual sperm might senesce during sexual rest/sperm storage. Sperm are metabolically active. Thus, sperm storage can rapidly lower sperm viability and motility due to the depletion of ATP reserves (Storey, 2008). For instance, guppy males with many days of sexual rest have less motile and slower sperm, as well as lower fertilisation success than recently mated males with fresh sperm (Cattelan and Gasparini, 2021; Gasparini et al, 2017; 2018). Similarly, queen ants lose viability of sperm stored over many years (Den Boer et al, 2009). Such declines are likely to reduce the fertilisation success of stored sperm. Mature sperm stored in-vivo also accumulate oxidative damage, which can lead to the fragmentation of sperm DNA (Barbagallo et al, 2022; Cattelan and Gasparini, 2021; Kugelman et al, 2024; Pizzari et al, 2008). Sperm lack cytoplasm and are mostly transcriptionally silent, resulting in insufficient DNA repair to remediate DNA fragmentation (Balder et al, 2024). For example, sperm storage during sexual rest increases sperm oxidative stress in insects (Reinhardt and Ribou, 2013) and sperm DNA fragmentation in men (Du et al, 2024; Barbagallo et al, 2022; Kugelman et al, 2024). Such intra-cellular declines in sperm can potentially impact embryos and offspring (reviewed in Barbagallo et al, 2023; Tarin, 2000). For instance, maternal (Jalme et al, 1994; Nalbandov and Card, 1943) and paternal (White et al, 2008) sperm storage in some birds reduces embryo survival, while paternal sperm storage in fish and fruit flies lowers the reproductive success of sons (Gasparini et al, 2017; Sanghvi et al, 2024a). Despite these and other studies showing that sperm storage negatively impacts sperm performance or offspring, some studies have not found such evidence (e.g. Firman et al, 2015; Meunier, et al, 2022; Vega-Trejo et al, 2019). This heterogeneity between study outcomes highlights the lack of consensus on whether or how sperm storage leads to post-meiotic sperm senescence, and our incomplete understanding of the biological and methodological factors that modulate these effects.

Several biological factors might influence the magnitude of sperm quality declines during sperm storage potentially explaining differences between study outcomes. The species-specific sperm-storing-sex, whether male or female, is likely one such factor. Males and females have evolved different mechanisms to maintain sperm quality and function during storage. Males in some species reabsorb (Lambiase and Amann, 1969) or even spontaneously discard old sperm (Quay, 1987), such as through masturbation (Brindle et al, 2023; Thomsen, 2000) and sperm dumping (Parnes et al, 2006; Reinhardt and Siva-Jothy, 2005). Furthermore, males produce seminal fluid with proteins and antioxidants (Fricke et al, 2023) that extend the longevity of stored sperm (Avila et al, 2011; Den Boer et al, 2008). Additionally, when sperm are not stratified by age and mix freely in male storage organs, constant spermatogenesis might prevent increases in average sperm age despite sperm being stored for extended durations (Reinhardt, 2007). Females, too, have strategies to potentially alleviate the effects of post-meiotic sperm senescence. Females can bias the storage of sperm and fertilisation of eggs towards younger sperm (e.g. Snook and Hosken, 2000; Vuarin et al, 2019). Moreover, in some species, female reproductive fluid and epithelial cells have protective functions that keep sperm viable for long durations (Baer et al, 2009; Gasparini and Evans, 2013; Holt and Fazeli, 2016) and increase the fertilisation success of aged sperm (Hadlow et al, 2023). These fluids might even explain why sperm stored in females have lower levels of oxidative stress than sperm stored in males (e.g. Ribou and Reinhardt, 2012). Females can also increase their rate of re-mating for fresh sperm when stored sperm accumulate higher levels of oxidative stress (Turnell and Reinhardt, 2022). Another potentially important biological modulator of post-meiotic sperm senescence is the specific trait measured (Reinhardt et al, 2015). Some sperm traits are likely to be more prone to temporal damage than others. For example, sperm storage might reduce sperm performance, therefore compromising traits related to fertilisation. However, the impact of sperm storage on offspring traits may be minimal if post-fertilisation mechanisms repair sperm damage, or if damaged sperm are selectively excluded from fertilisation (see Sanghvi et al, 2024b, who show this for male age).

In addition to biological variables, methodological factors could potentially modulate the effects of sperm storage. For example, differences in the sampling method used to assay individuals could lead to differences between study outcomes. Longitudinal sampling of the same individual across different sperm storage durations minimizes the impact of genetic differences between individuals, and accounts for the confounded influence of other variables, such as age in species where sperm storage represents a small fraction of lifespan. Thus, longitudinal sampling can potentially lead to a higher detectability of post-meiotic sperm senescence compared to cross-sectional sampling (Sanghvi et al, 2024b). If sperm quality decline during storage is linear, then the exact duration for which sperm are stored might also modulate storage effects such that longer sperm storage durations lead to more deterioration in sperm and offspring quality.

Assessing the overall extent of sperm senescence during sperm storage requires an integrative analysis that includes data from various taxa and both sexes, as well as theory from biomedical and evolutionary fields while considering the specific traits and sampling methods. Here, we conduct a meta-analysis across humans and non-human animals to understand the overall outcomes of in-vivo sperm storage on sperm health and fitness. Specifically, we address three aims: (i) document effects of in-vivo sperm storage on sperm quality, fertilisation, and post-fertilisation outcomes across animals; (ii) assess the impact of different moderating variables such as the sex of the sperm-storing individual, type of sampling, and sperm storage duration; (iii) compare the impacts of sperm storage on traits related to sperm health and performance versus male fitness.

## Methods

We conducted and reported our meta-analysis following the PRISMA-EcoEvo guidelines (O’Dea et al. 2021), and performed all statistical analyses using R version 4.1.2 (R Core Team, 2020). We searched, screened, and analysed studies on humans versus non-human animals (henceforth, “animals”) separately, given the stark biological and methodological differences between the two types of studies (see Discussion), and because there were substantially more studies in humans than in animals on the topic. Our pre-registration report, data set, and code can be found at OSF (10.17605/OSF.IO/JXV7Z).

### Search Protocol

We generated appropriate search strings to find relevant papers separately for human and animal studies, based on commonly used words found in the abstracts and keywords of previous studies on this topic (Appendix 1, Figure S1 for search string). For humans, we searched for relevant studies using SCOPUS and the Web of Science on January 26, 2024. In addition, we performed backward and forward searches for papers using six systematic reviews on male sexual abstinence (Appendix 2) and identified two further studies from other sources. To obtain relevant papers on animals, we conducted a literature search using SCOPUS and Web of Science on September 21 and October 12, 2022, respectively, and a backward search using 11 relevant reviews (Appendix 2). For animals, we additionally included eight studies identified through other sources, such as Google Scholar or those found in the human dataset. Two animal studies were from unpublished but publicly available Ph.D. theses. After removing duplicates, we obtained 1,069 on humans and 7,559 studies on animals. The abstracts of these studies were screened to assess their relevance for our meta-analysis.

### Screening Criteria

Abstracts for both human and animal studies were evaluated using predefined inclusion and exclusion criteria, in the software Rayyan (Ouzzani et al, 2016). Studies were excluded if they: addressed irrelevant topics, only reported the presence or absence of sperm storage, focused solely on the descriptive biology of spermatogenesis or sperm storage organs, were case studies with a sample size of one, involved plants (e.g. on endosperm, meiosis in sperm), had overlapping sperm storage durations between groups, were not written in English, compared sperm storage across species rather than within-species, were reviews or meta-analyses. This initial screening identified 315 relevant studies on humans and 177 on animals. These were then subjected to full-text screening to confirm their suitability.

At the full-text screening stage, we applied additional exclusion criteria (Appendix 1) while also reapplying all criteria from the abstract screening stage. This ensured the removal of any studies that may have initially passed abstract screening due to insufficient detail but were later found to be irrelevant upon full-text review. We excluded studies that: stored/aged sperm in-vitro or ex-vivo (i.e. outside the live individuals via cryopreservation or in a medium, e.g. Levitan, 2000); addressed unrelated topics (e.g. sperm precedence or alcohol abstinence instead of sexual abstinence); were inaccessible; did not include variation in, or manipulation of, sperm storage durations; had unclear experimental designs; measured traits without a clear relation to sperm health and performance or male fitness outcomes (e.g. offspring sex ratio); lacked sufficient data for calculating effect sizes (e.g. missing sample sizes). Crucially, we also excluded studies where outcomes could be attributed directly to variation in sperm quantity or to ejaculate depletion, because our focus was on post-meiotic sperm senescence which is the change in sperm quality due to storage (Pizzari et al, 2008; Reinhardt, 2007). For instance, traits like the number or proportion of eggs fertilised were excluded (e.g. Jones and Elgar, 2004) unless sperm numbers available during fertilisation were same across storage treatments. This is because individuals sampled at different sperm storage timepoints would have different numbers of sperm available for fertilisation, altering fertilisation rates due to sperm depletion or accumulation rather than just post-meiotic sperm ageing. Similarly, studies on competitive fertilisation success were excluded unless equal ejaculate quantities were used for insemination or were available for fertilisation between different sperm storage treatments.

### Data Collection

To investigate the effects of in-vivo sperm storage on sperm traits, fertilisation success, and post-fertilisation outcomes, we collected data on the means, standard deviations (SD) or standard errors (SE), as well as sample sizes of individuals for each sperm storage group/treatment (Appendix 3). We additionally recorded the number of unique and total individuals sampled per study. Total individuals sampled exceeded unique number of sampled in longitudinal (i.e. when individuals were sampled repeatedly at different durations of sperm storage/sexual rest) and semi-longitudinal studies (some individuals were repeatedly sampled while others not). In human studies, all data focused on male sperm storage due to a lack of studies on female storage, thus, the numbers of males sampled was recorded as our sample size. However, in animal studies, both males or females were sperm-storing individuals, and here, the sample size recorded corresponded to the numbers of individuals of the sex that stored sperm. When means and SD/SE were not reported, we extracted test statistics (e.g. t, F, Chi sq.-values or R²) to compute effect sizes (more below, formulae from Koricheva et al, 2013; Nakagawa and Cuthill, 2007; Polanin and Stiltsveit, 2016). To ensure accuracy of the data collection procedure, we verified the repeatability of data extraction for both human and animal datasets, and these were highly repeatable (Appendix 4).

To examine how moderators influenced the effects of sperm storage, we collected data on various biological and methodological variables from included studies. For human studies, we collected data on country, trait (Appendix 5), health condition of males (fertile, infertile-i.e. attended infertility clinics, or undergoing treatment for other conditions), sperm storage duration range, and male sampling method. For animal studies, our moderators included: species; taxonomic class; trait (Appendix 5); range of sperm storage duration (i.e. difference between the maximum and minimum storage duration in years for each effect size); sex of the sperm-storing individual; male sampling method (longitudinal, semi-longitudinal, or cross-sectional); and population type (laboratory, captive, wild, or domestic). We furthermore collected data on the maximum lifespans of included species from comprehensive animal databases (e.g. Animal Diversity Web and AnAge-DeMagalhaes and Costa, 2009), a previous meta-analysis on male ejaculate ageing (Sanghvi et al, 2024b), or primary species-specific literature. This was done to additionally standardise sperm storage durations proportional to the species’ maximum lifespans allowing comparisons across species using absolute as well as relativised durations of sperm storage. Data on sperm storage durations for both humans and animals were log_e_ transformed for normality because the original distribution was skewed (Appendix 6).

Overall, our meta-analysis contained data on traits related to sperm performance (i.e. motility, velocity, viability, morphology, fertilisation success), sperm intra-cellular quality (i.e. oxidative stress and DNA quality), and post-fertilisation outcomes (i.e. embryo or offspring quality).

### Effect Size Calculation

We first calculated Pearson’s correlation coefficients using four different methods: (i) standardised mean differences for comparisons of two storage groups with reported SD/SE (Appendix 7A); (ii) simulations for multiple storage duration group comparisons with reported means, SD/SE (Appendix 7B); (iii) test statistics (e.g., t-values, Z-values) (Appendix 7C); (iv) percent/proportion data when SD or SE was unavailable (Appendix 7D). We then calculated Fisher’s z-transformed correlation coefficient as our effect size from the calculated Pearson’s correlation coefficients (Nakagawa and Cuthill, 2007; Appendix 7E). Effect sizes were standardised using a multiplier to ensure that negative values indicated trait deterioration (and potentially lower fitness) with sperm storage, while positive values indicated improvement (and potentially higher fitness).

Sampling variance of effect sizes was calculated as: 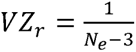 where *N_e_* was the effective sample size (Appendix 8). For longitudinal and semi-longitudinal studies, the effective sample size accounted for pseudo-replication of individuals sampled repeatedly at different time points following Rutkowska et al (2014). For cross-sectional studies, total individuals, which equalled number of unique individuals, was the effective sample size.

### Data Analysis

We analysed human and animal datasets separately. For both datasets, we first constructed a meta-analytical model to assess the overall effect of sperm storage on traits using the rma.mv function in the *metafor* package (Viechtbauer, 2010). In all our models, effect size (Zr) was the response variable. We computed a variance-covariance (VCV) matrix with the *vcalc* function to estimate effect size variances, containing VZr (diagonal) and within-cohort covariances (off-diagonal, rho = 0.8), and used this VCV matrix as the sampling variance (Nakagawa et al, 2023; Noble et al, 2017). Random effects included in both, animal and human models for the VCV matrix were: effect size ID (residual variance), cohort ID (within-study variance), and study ID (between-study variance). For human data, we also included country of origin of the studied men as a random effect to address possible non-independence caused by geographic and cultural history. On the other hand, for animal data, we included species ID and a correlation matrix of phylogenetic relatedness as random effects. The phylogenetic tree was constructed with the *ape* (Paradis et al, 2004) and *rotl* (Michonneau et al, 2016) packages which use relatedness data from OpenTreeOfLife. Total heterogeneity (I²) and partial heterogeneity for random effects were estimated using the i2_ml function in the *orchard* package (Nakagawa et al, 2023).

Next, we constructed meta-regressions to explore how moderators influenced the effects of sperm storage. “Full models” were constructed for each dataset to include all moderators relevant to that dataset: trait, health condition, sperm storage duration (log_e_-transformed), and sampling method for humans; trait, sex of storage, taxonomic class, male sampling method, sperm storage duration (log_e_-transformed absolute and standardised), and population type for animals. Full models assessed the proportion of heterogeneity explained by each moderator while controlling for confounding effects of other moderators (Noble et al, 2022). We additionally created individual meta-regressions separately for each moderator to test the influence of individual variables on modulating the influence of sperm storage. Models with intercepts were used to test whether the variance explained by moderators was significant. Models without intercepts were used to quantify whether each level of a moderator was significantly different from zero (i.e. showing deterioration or improvement). For all our meta-regression models, we calculated total heterogeneity (Q_M_) and marginal R² for moderators (significance of α= 0.05).

Note: Effect sizes in Results are presented as Pearson’s correlation coefficients back transformed from Zr for ease of interpretation.

### Publication Bias

We assessed publication bias for human and animal datasets separately using the following methods (Nakagawa et al, 2022): (i) funnel plots, to visually assess whether less precise studies with non-significant results were missing as indicated by asymmetry in effect size distributions; (ii) trim-and-fill analysis, with effect sizes aggregated at the study-level, to quantify missing studies; (iii) time lag bias analysis, to test whether year of publication influenced effect sizes; (iv) small-study bias analysis, to test whether the sample size of a study influenced the magnitude of effect sizes; (v) selection model, to test whether studies with larger effect sizes were more likely to be included.

## Results

### Dataset description

The human dataset contained 453 effect sizes from 115 studies, representing at least 54889 unique men from 31 countries. The animal dataset contained 235 effect sizes from 56 studies representing 30 species (Appendix 1). Both human and animal studies were each equally represented across cross-sectional and longitudinal sampling. There were more human studies representing sperm performance than post-fertilisation traits, and more animal studies representing sperm performance than sperm intra-cellular traits. There was collinearity between male sampling and method of effect size calculation in humans (Figure S4A), and between the sex of storage and type of trait in animals (Figure S4B). The duration (mean and variance) of sperm storage was higher in animals than humans (Figure S2, S3).

### Humans

Sperm storage negatively impacted outcomes in men (Pearson’s *r* [95% C.I] = -0.093 [-0.120 to -0.066], Figure 1A). There was significant heterogeneity in our data (I^2^ = 97.593, P < 0.001), all of which was attributed to differences between effect sizes (i.e. residual variance), with 0% of heterogeneity being explained by true difference between studies (Figure S5), country, or cohort. Including all moderators in our meta-analytical “full” model collectively explained a significant proportion of variation (R^2^ = 12.693%, Q_M_ = 40.691, Q_E_ = 30060.027, P < 0.001, DF_11,431_).

**Figure 1:**
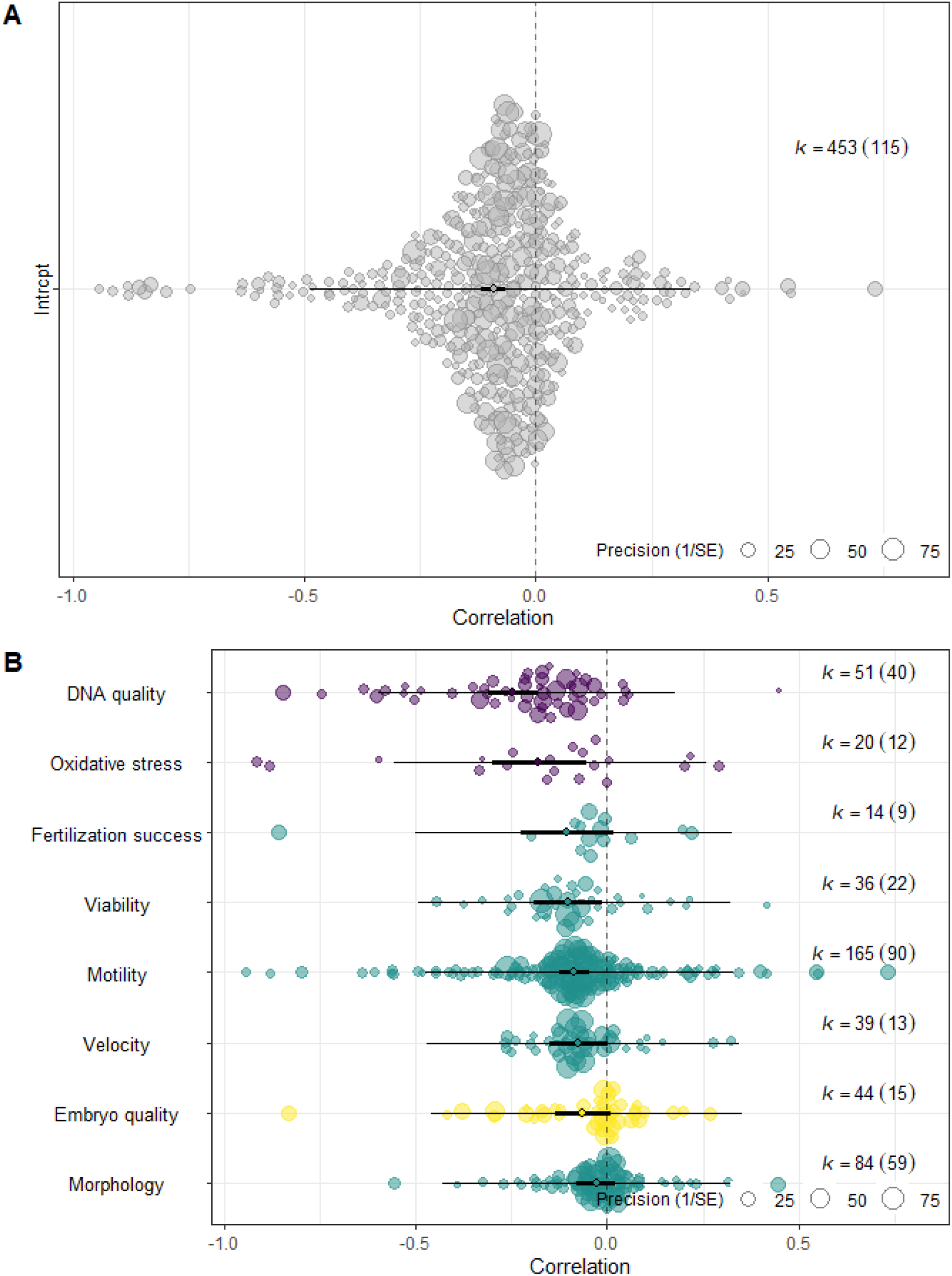
**(A)** Null model shows overall deleterious effects of sperm storage in humans (i.e. men). **(B)** Consequences of sperm storage for different sperm performance-related (green), sperm intra-cellular quality (purple), and post-fertilisation (yellow) traits in humans. Negative correlations (Zr back-transformed to Pearson’s r) depict poorer reproductive outcomes when sperm are stored, and positive values represent improvements in reproductive outcomes with storage. The size of each datapoint represents precision of the effect size. Bold error bars show confidence intervals, and light bars show prediction intervals. Samples sizes k = number of effect sizes (number of studies in brackets).

Trait-specific impacts of sperm storage explained a significant amount of heterogeneity (Q_M_ = 30.182, Q_E_ = 30524.926, DF_7,445_, R^2^ = 7.956%, P < 0.001). Specifically, sperm motility and viability (i.e. sperm performance) significantly decreased, and sperm DNA damage and oxidative stress (i.e. intra-cellular sperm quality) significantly increased during storage (Figure 1B). Embryo quality, fertilisation rate, and sperm velocity were marginally impacted (P < 0.092), while sperm morphology was not significantly impacted (Figure 1B). Studies that sampled men longitudinally found more severe effects of sperm storage than cross-sectional studies (Figure 2A), and this difference explained significant heterogeneity in the data (Q_M_ = 12.389, Q_E_ = 34544.397, DF_1,451_, R^2^ = 5.649%, P < 0.001). Neither the specific range of time for which men were sexually abstinent in a study (Q_M_ = 0.007, R^2^ = 0; P = 0.934, Figure 2B), nor the health condition of males (R^2^ = 0%, P = 0.897, Figure S6) explained significant heterogeneity.

**Figure 2:**
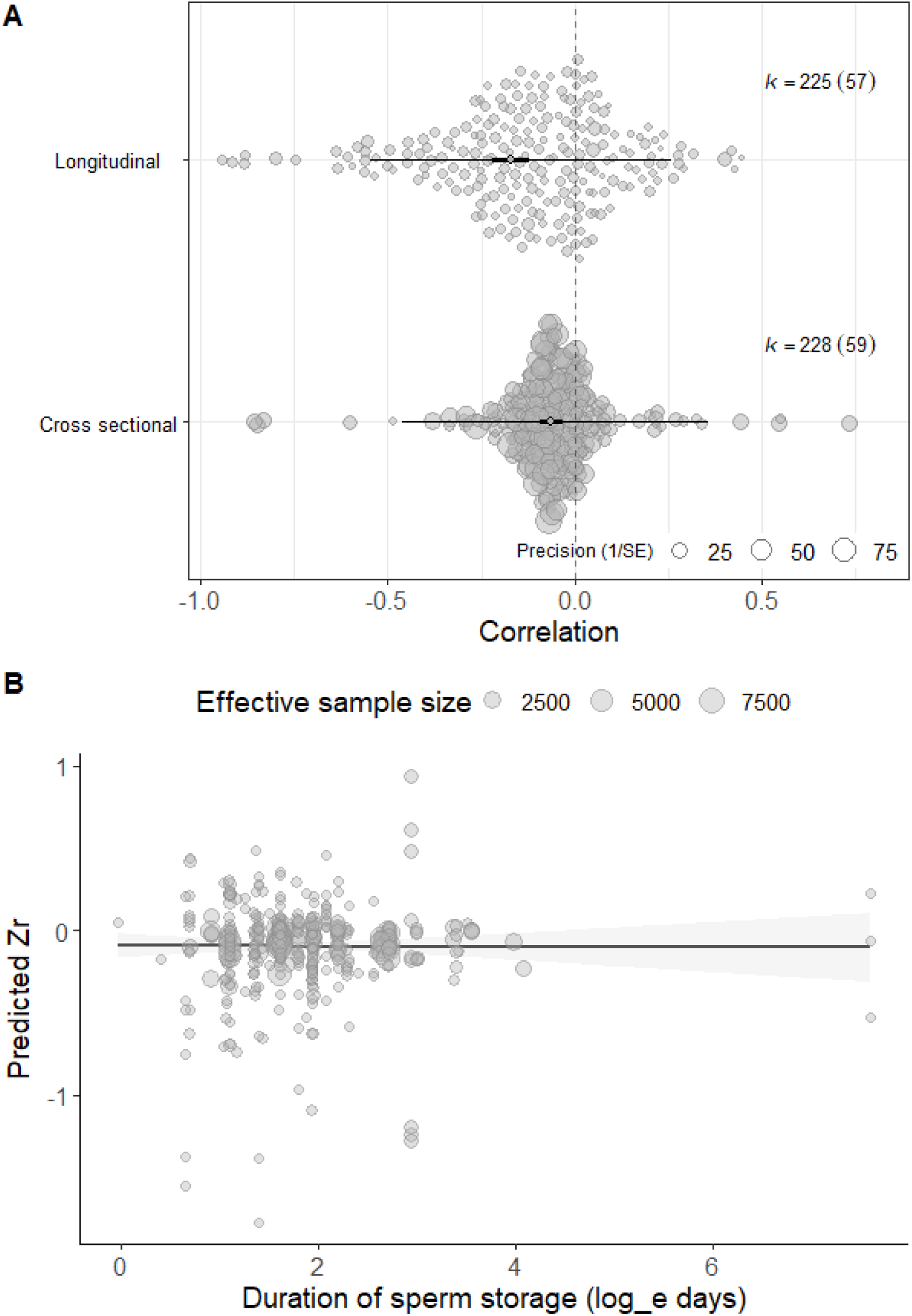
**(A)** Sampling method in humans significantly modulated the impact of sperm storage on outcomes. Negative correlations (Zr back-transformed to Pearson’s r) depict poorer reproductive outcomes when sperm are stored, and positive values represent improvements in reproductive outcomes with storage. Bold error bars show confidence intervals, and light bars show prediction intervals. k = number of effect sizes (number of studies). **(B)** Range of sampling duration of sperm (log_e_) storage/sexual abstinence did not significantly impact outcomes (Zr) in humans. Size of points represents effective sample size. 1 day = 0 log_e_ days, 10 days = 2.3 log_e_ days, 50 days = 3.9 log_e_ days

### Animals

Heterogeneity among effect sizes in animal studies was high (I^2^ = 82.782%, P < 0.001) with 43.104% attributed to true differences between studies (Figure S7), 4.513% to between-species differences, but 0% to between-cohort differences or phylogenetic relatedness (Figure 3A). Like in humans, sperm storage led to significant deterioration in recorded traits in other animals (Pearson’s *r* [95% confidence interval (C.I.)]: -0.198 [-0.293 to -0.102], z = -4.057, P < 0.001, Fig. 3B). All included moderators in our meta-regression model collectively explained substantial heterogeneity (R^2^ = 22.274%) in the data, although this was non-significant likely due to the low sample size to moderators ratio (Q_M_ = 22.775, Q_E_ = 1194.243, P = 0.089, DF_15,214_).

**Figure 3.**
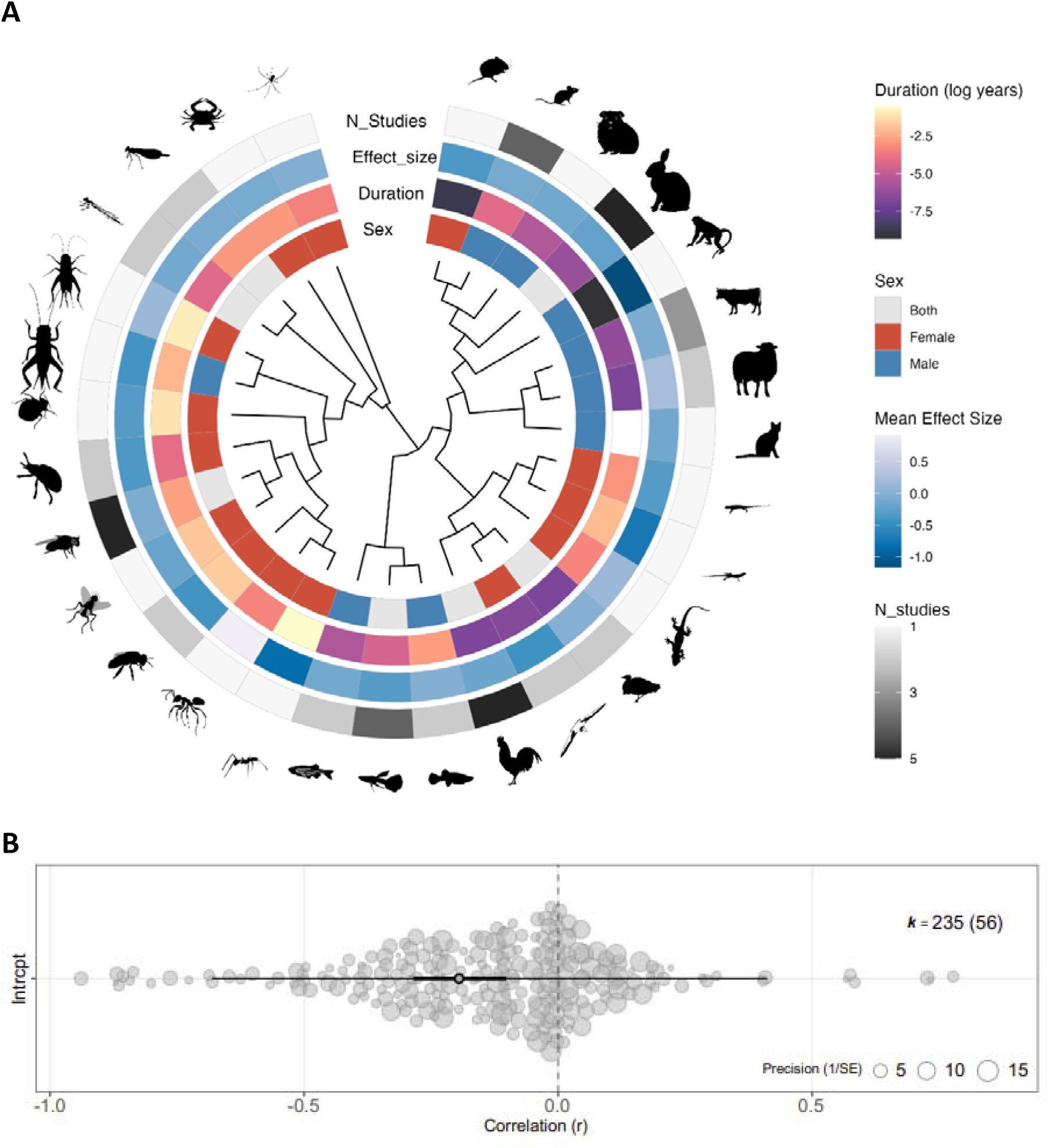
**(A)** Phylogenetic tree of species included in the animal dataset, specifying the sex sperm storage takes place within, the duration of sperm storage (in log_e_ years), the mean effect size for each species (Zr back-transformed to Pearson’s r), and the number of studies. 1 hour = -9.079 log_e_ years, 1 day = -5.9 log_e_ years, 1 week = -3.96 log_e_ years, 1 month = -2.485 log_e_ years, 1 year = 0 log_e_ years. **(B)** Null model shows overall effects of sperm storage in animals. Negative correlations (r) depict poorer reproductive outcomes when sperm are stored, and positive values represent improvements in reproductive outcomes with storage. The size of each datapoint represents precision. Bold error bars show confidence intervals, and light bars show prediction intervals. k = number of effect sizes (number of studies). Silhouettes representing included species taken from Phylopics under CC0, CC-BY, CC1.0 licences.

Despite the mean effect size being more negative for females than males, this difference was not significant (Q_M_ = 1.355, Q_E_ = 1347.467, P = 0.246, R^2^ = 2.365%), and both male (P = 0.035) and female (P < 0.001) sperm storage reduced fitness outcomes (Figure 4A). The impact of sperm storage tended (non-significantly) to depend on the measured trait (Q_M_ = 13.976, Q_E_ = 1280.619, DF_8,226_, R^2^ = 6.429%; P = 0.082, Figure 4B, Figure S8A). Sperm viability, velocity, motility, morphology, fertilisation success (all measures of sperm performance), and embryo quality (a measure of post-fertilisation outcomes) were impacted negatively (Figure 4B). Sperm DNA damage or oxidative stress, (both measures of intra-cellular sperm quality) and offspring quality were not significantly influenced (Figure 4B). The specific range of absolute and standardised durations for which sperm were stored significantly impacted measured outcomes 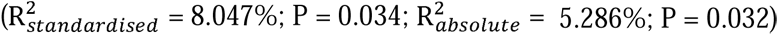. Specifically, longer durations of storage led to more severe deterioration in measured outcomes and this pattern was more severe for male than female sperm storage (Figure 5). Population setting did not significant modulate outcomes despite lab animals experiencing more negative outcomes than wild, captive, or domestic animals (R^2^ = 4.729 %, P = 0.573, Figure S8B). Taxonomic class (R^2^ = 0.741%, P = 0.996, Figure S8C) or sampling method (R^2^ = 0.357%, P = 0.652, Figure S8D) did not explain significant heterogeneity.

**Figure 4.**
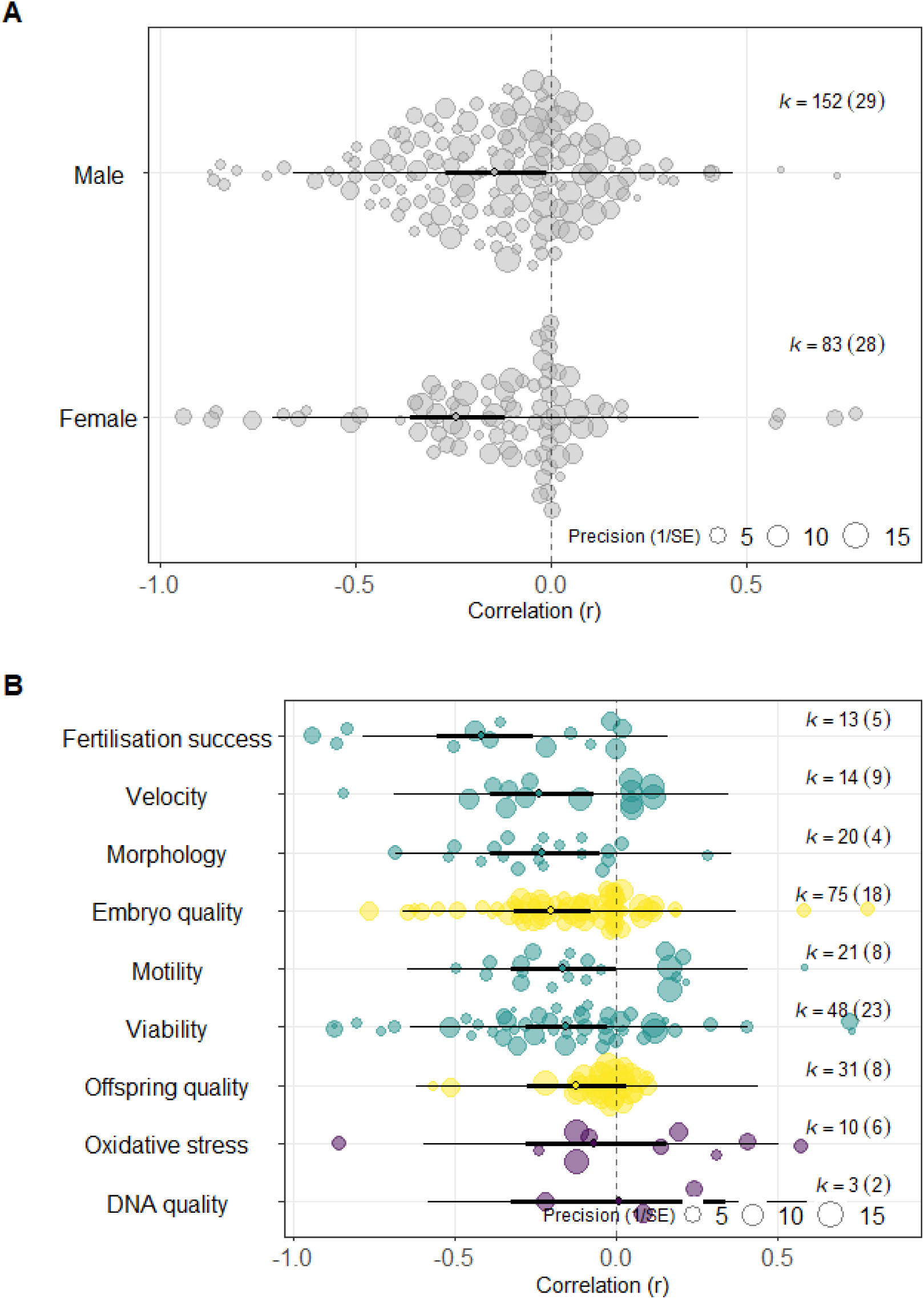
**(A)** Overall effect of sperm storage in male and female animals. (**B)** Consequences of sperm storage for different sperm performance-related (green), intra-cellular quality (purple), and post-fertilisation (yellow) traits. Negative correlations (Zr back-transformed to Pearson’s r) depict poorer reproductive outcomes when sperm are stored, and positive values represent improvements in reproductive outcomes with storage. The size of each datapoint represents precision. Bold error bars show confidence intervals, and light bars show prediction intervals. k = number of effect sizes (number of studies).

**Figure 5.**
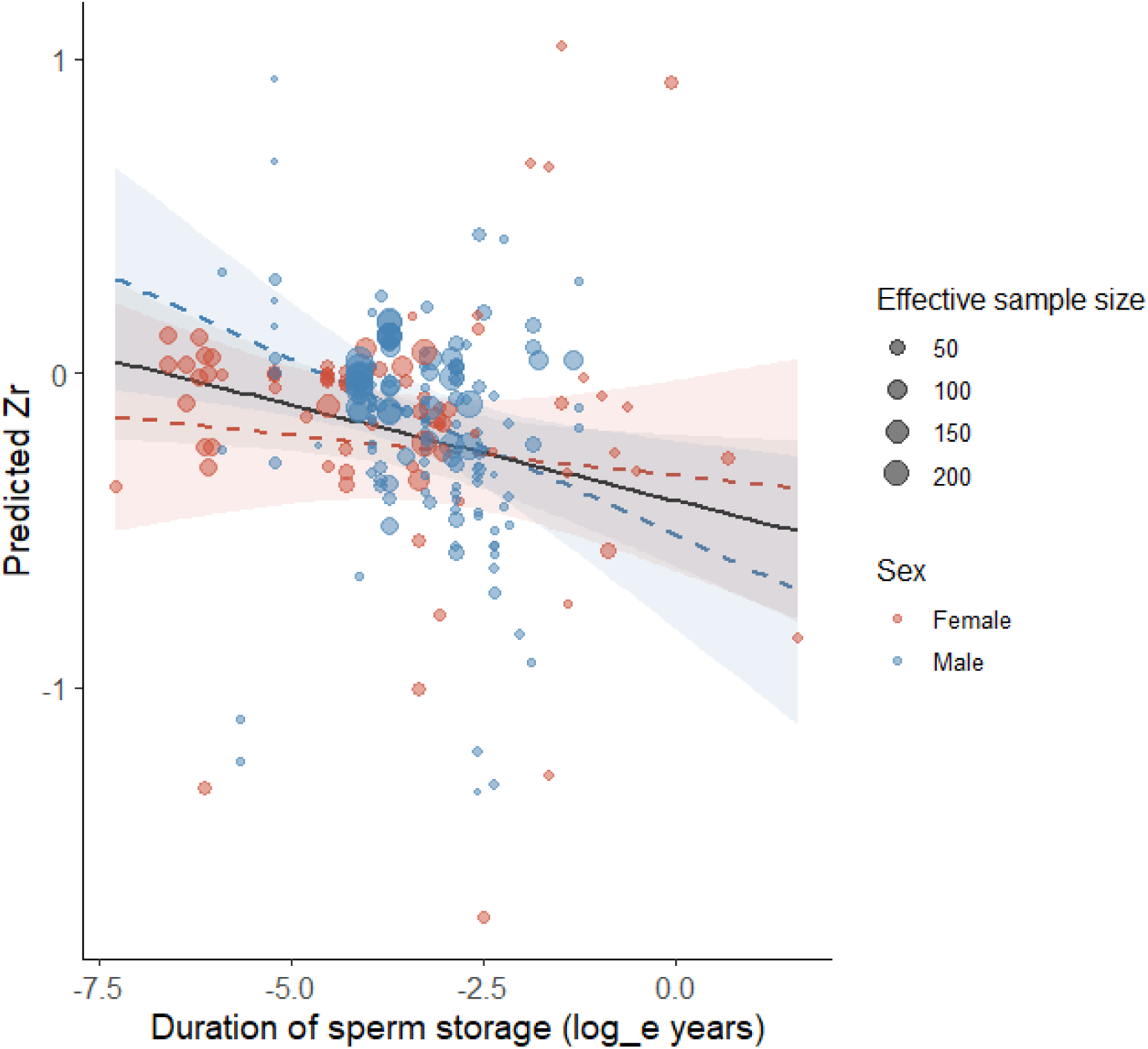
Range of sperm storage duration sampled by a study negatively impacts reproductive outcomes (Zr). Females have more negative intercepts but shallower slopes than males. Size of data point reflects the effective sample size. 1 day = -5.9 log_e_ years, 1 week = -3.96 log_e_ years, 1 month = - 2.485 log_e_ years, 1 year = 0 log_e_ years.

### Publication bias

We found no statistical evidence of publication bias in our human dataset (Figure S9) or animal dataset (Figure S10), as assessed by: visually checking funnel plots; constructing an aggregated trim and fill model (humans: estimated missing studies = 0, SE = 5.459; animals: estimated missing studies = 0, SE = 3.994); assessing small study bias (humans: t = -1.492, P = 0.136; animals: t = -0.820, P = 0.413) and time lag bias (humans: t = -0.617, P = 0.548; animals: t = 0.498, P = 0.619). Selection model (humans and animals: P = 0.001), surprisingly showed larger effect-sizes having a lower probability of being selected.

## Discussion

In-vivo sperm storage in males and females has been observed across the animal kingdom. Despite such expected ubiquity, surprisingly little quantitative synthesis exists on how sperm storage impacts sperm quality, fertilisation outcomes, and offspring phenotypes, thus consequently, organismal fitness. Our meta-analysis explores the overall taxon-wide evidence for whether sperm storage leads to post-meiotic sperm senescence (Pizzari et al, 2008; Reinhardt, 2007). In general, we find considerable evidence for deterioration in sperm performance traits, and to some extent, fertilisation success and post-fertilisation outcomes. We discuss the implications of sperm deterioration during sperm storage for male and female fitness outcomes and explore the impact of biological and methodological moderators on sperm storage.

### Mechanisms of trait-specific effects

Post-meiotic sperm senescence during sperm storage is hypothesized to occur through two primary mechanisms: reduction in sperm viability or performance due to depletion of ATP reserves (Den Boer et al, 2009; Storey, 2008); and reactive oxygen species causing irreparable DNA damage (Barbagallo et al, 2022; Cattelan and Gasparini, 2021; Kugelman et al, 2024; Pizzari et al, 2008) and ultimately male infertility (Agarwal and Said, 2003; Sharma and Agarwal, 1996). The former mechanism is likely to impact fertilisation success of individual sperm, while the latter mechanism could potentially have inter-generational impacts on embryo and offspring quality (Barbagallo et al, 2022; Tarin, 2000). In humans, we found support for both mechanisms driving overall deleterious outcomes. In other animals, sperm storage reduced sperm performance and embryo quality, but did not impact intra-cellular traits of DNA or oxidative damage, which could be due to small sample sizes in animals for these traits. Despite this, we found negative effects on embryo quality indicating potential for extended sperm storage being selected against. Future studies could investigate whether changes in sperm gene expression due to epigenetic modifications during sperm storage might also lead to inter-generational effects (Chen et al, 2016).

One reason for differential impacts of sperm storage on different traits could be due to variation in the likelihood, onset, shape, and rate of deterioration in each trait/outcome (Reinhardt, 2007; Reinhardt et al, 2015; Sanghvi et al, 2024b), akin to senescence in whole organisms (Jones et al, 2014; Lemaitre and Gaillard, 2017), for which there could be several mechanisms. For example, if within a stored ejaculate, deteriorated sperm are less likely to fertilise eggs (Cattelan and Gasparini, 2021; Gasparini et al, 2017; 2018), then post-meiotic sperm senescence in sperm performance is unlikely to impact post-fertilisation outcomes. Similarly, if females or embryos can repair damage in the DNA of stored sperm after fertilisation (reviewed in Garcia-Rodriguez et al, 2018; Martin et al, 2019; Newman et al, 2022), then deterioration in intra-cellular sperm traits is unlikely to impact embryo or offspring quality. A meta-analysis on male ageing by Sanghvi et al (2024b) found such results, whereby ejaculate traits were more likely to decline with male age compared to fertilisation or post-fertilisation traits. Our analyses generally suggest that despite sperm storage reducing sperm performance (i.e. motility, velocity, viability, fertilisation success), post-fertilisation outcomes might be more protected.

### Humans versus animals, males versus females

While humans and animals showed similar overall outcomes due to sperm storage, they differed in the specific traits that were impacted and the moderators that explained substantial variance. This might be due to smaller sample sizes in animal studies, which reduce statistical power. However, differences in selection pressures, environments, and life-history strategies between humans and other animals could cause variation in resource allocation towards different sperm traits and also explain our results. Alternatively, these discrepancies could be due to differences between how human versus animal studies were conducted. For instance, human studies included individuals who were both infertile as well as healthy whereas animal studies did not have such distinctions. Additionally, human studies focussed on male sperm storage while animal studies included both male and female storage (e.g. Wetzker et al, 2024).

The duration of abstinence or sperm storage need not reflect the precise post-meiotic age of the sperm (Box 1). For example, in our human’s dataset, longitudinal studies often collected a first ejaculate from men after prolonged periods of sexual abstinence, and then a second ejaculate after a shorter duration of abstinence (e.g. Agarwal et al. 2016; Gonsalvez et al. 2011, Box 1). Here, the second ejaculate would on average have been stored for a shorter duration and contain more fresh sperm than the first. However, because men were unlikely to be completely sperm depleted after their first collection, the actual duration of shorter sexual abstinence would not precisely equate to the age of sperm in the second ejaculate (Figure 6A). Even in animals, when studies "stripped" males of their stored ejaculates completely before sampling sperm (e.g. Gasparini et al, 2018), ongoing spermatogenesis and sperm reabsorption could have altered the ages of individual sperm in the stored ejaculate. Thus, even under such stripping conditions, the duration of storage/sexual rest in men would only represent the maximum post-meiotic age of the sperm. However, in female animals, the duration for which sperm were stored before being used to fertilise eggs would accurately correspond to the minimum post-meiotic ages of sperm. Future studies could test how differences in the timing of sperm inseminated in women prior to ovulation impacts fertilisation rates and embryos. Given our results in sperm, we also suggest that studies could quantify whether storage of oocytes/eggs might impact egg and offspring quality (reviewed in Tarin et al, 2000) above and beyond the confounded influence of maternal age (Woodruff, 2008).

**Figure 6:**
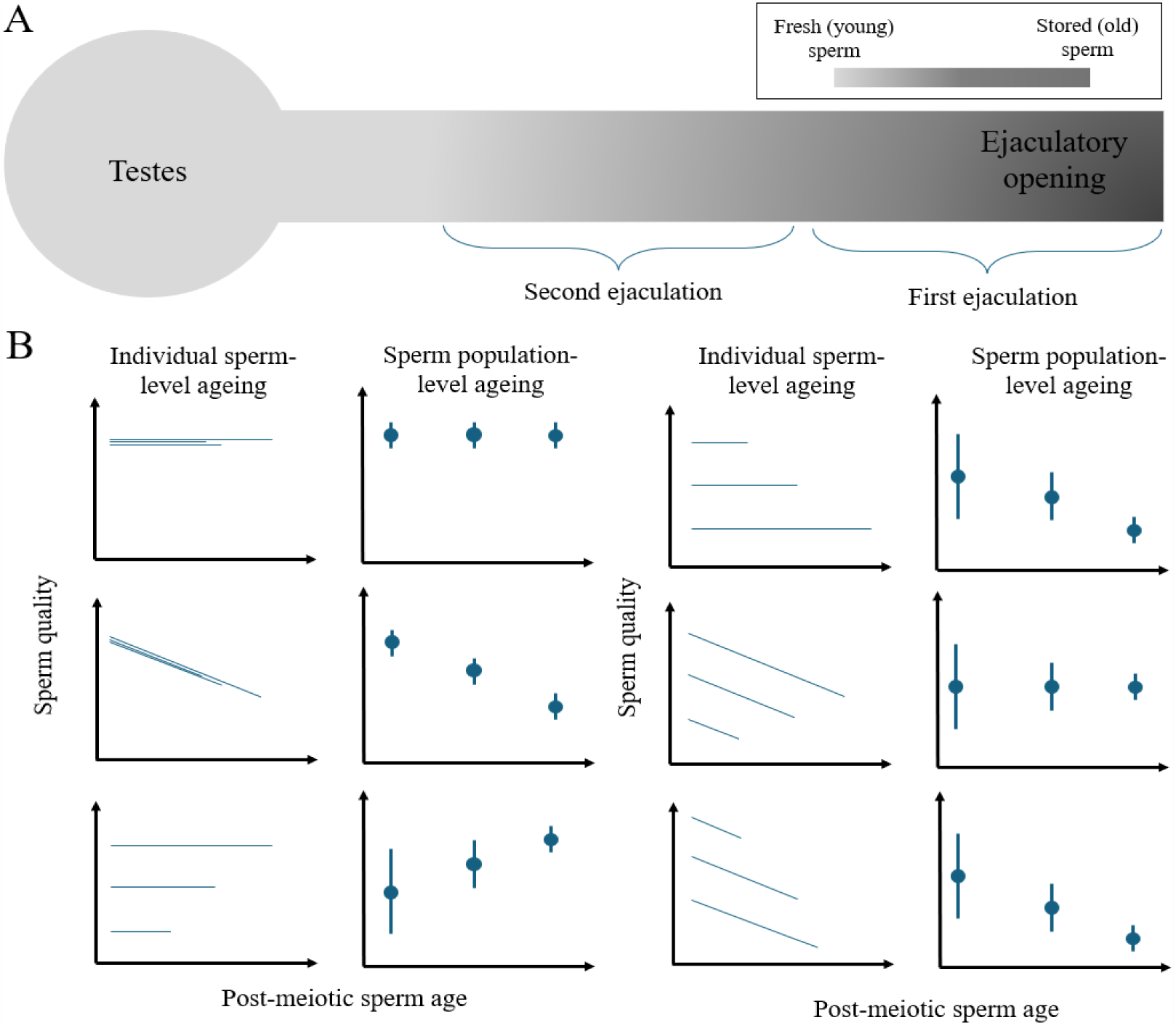
**(A)** Sperm stratification results in older, more deteriorated sperm being closer to the ejaculatory opening. Thus, in longitudinal studies, the first ejaculation contains older sperm, regardless of sexual rest duration. The second ejaculation contains younger sperm, however, the duration of sexual rest when obtaining the second ejaculate does not correspond to the exact age of the sperm unless males have been fully “stripped” first. In contrast, sperm mixing and continuous replenishment prevent mean sperm age increases. **(B)** Effects of sperm ageing on quality depend on between-sperm variation (e.g. due to selective disappearance) and within-sperm changes (e.g. senescence in quality). Each lines represents an individual sperm; dots and error bars show ejaculate-level means and variance in sperm quality. Termination of a line represents death/resorption of sperm.

#### Box 1

**Storage does not imply post-meiotic sperm senescence**

Despite individual sperm advancing in age during storage, the average age of ejaculates might not increase in males and sperm may not deteriorate, for various reasons. For example, if sperm are stratified such that freshly made sperm are close to the testes (distal) and older sperm close to the site of ejaculation (caudal) (Eckel et al, 2017; Squires et al, 2015), then increasing duration of sexual abstinence would lead to ejaculated sperm being older despite fresh sperm being produced (Kang et al, 2023; Reinhardt, 2007; Figure 6A). However, if sperm mixing occurs, then continuous production of fresh sperm (e.g. Sepil et al, 2020) or re-absorption of old sperm (Jones, 2004) could prevent the average age of the ejaculate from increasing. Additionally, the advancement of ejaculate age might not necessarily lead to deterioration (i.e. post-meiotic senescence) in sperm quality due to selective disappearance of sperm (Van de Pol and Verhulst, 2008) caused by between-sperm variation and non-random sperm death. Here, if sperm populations sampled at older post-meiotic ages represent a biased cohort of sperm that are better in quality (e.g. Alavioon et al, 2017, 2019; Crean et al, 2010; Marcu et al, 2024), then within-sperm post-meiotic senescence would be masked by sperm-population-level improvements (Figure 6B).

### Evolutionary consequences

Sperm storage in males and females for extended durations has evolved across the animal kingdom (Orr and Brennan, 2015; Pizzari et al, 2008; Reinhardt, 2007). The ability to store sperm for extensive durations likely provides an evolutionary advantage because it allows the temporal separation of spermatogenesis, ejaculation, and fertilisation, and allows sperm accumulation (Birkhead and Moller, 1993). However, given the costs of storage due to post-meiotic sperm senescence as demonstrated by our results, it is likely that animals have evolved various strategies to alleviate these effects. For example, sperm DNA have their histones replaced with protamine to tightly bind chromatin, thus reducing the amount of DNA exposed to reactive oxygen species (Agarwal, 2003; Drevet et al, 2022; Hamilton and Assumpcao, 2020). Additionally, in some species, both males and females spontaneously eject old sperm (Parnes et al, 2006; Snook and Hosken, 2004; Thomsen, 2000). An extreme example of this in males is masturbation, which has been hypothesized to have evolved as an adaptation for ejecting stored, old sperm (Thomsen, 2001; reviewed in Brindle et al, 2023) especially in species where sperm are stratified by age. Male production of seminal plasma/seminal fluid with antioxidant properties (Aitken et al., 2015; Peeker et al., 1997; Fricke et al., 2023) might also represent an evolutionary adaptation to mitigate sperm senescence. Similarly, in addition to specialised sperm storage organs in females (Holt and Fazeli, 2016), female reproductive fluid/ovarian fluid could have evolved to nourish sperm and extend their longevity (Baer et al, 2009; Gasparini and Evans, 2013).

Post-meiotic sperm senescence is likely to have important consequences for sexual selection and our understanding of polyandry, cryptic female choice, and last-male sperm precedence. For example, females might temporally partition sperm from different males in their storage organs (e.g. Birkhead and Hunter, 1990; Briskie, 1996; Carleial et al, 2020; King et al, 2002; Roderick et al, 2003) to preferentially use fresh sperm (e.g. Snook and Hosken, 2004). Females might also remate with the same male for younger sperm when sperm are stratified in males, leading to the first ejaculation containing older sperm. Females could also bias paternity towards last-mated males due to sperm from first-mated males having been stored for extended durations (Laturney et al, 2018). These predictions could be tested by future studies.

Our meta-analysis highlights the importance of understanding variation between sperm not only at the male-level, but also within-ejaculates and temporally within-sperm. This outlook places emphasis on the typically overlooked processes of haploid selection occurring at the within-ejaculate, between-sperm level. Recent evidence shows that selection can act on such temporally induced between- and within-sperm variation, having crucial implications for evolution and reproductive health (Alavioon et al, 2017, 2019; Crean et al, 2010; Marcu et al, 2024).

### Biomedical consequences

Results from our study have important consequences for human and animal fertility, such as artificial insemination in human assisted reproduction techniques like IVF or ICSI and captive breeding programs. Garcia-Grau et al (2022) have suggested that the currently declining fertility rates across some countries might be due to negative effects of sexual abstinence on sperm quality. Sokol et al (2021) recommend that the present WHO guidelines, which suggest a 2–7-day abstinence period, might be too long if sperm quality is to be optimised. Our results generally support the conclusion that within a study (but not across studies-see Figure 6A for why), longer periods of sexual abstinence/sperm storage are deleterious for sperm quality. We suggest that future studies investigate whether variation in abstinence durations across past decades might explain the observed declines in fertility of men over the past years.

Despite the negative effects on sperm quality, sexual rest across animals leads to accumulation of sperm numbers (Du et al, 2024; Hanson et al, 2018; Sanghvi et al, 2024b; Sanghvi et al, 2025; Sepil et al, 2020). Therefore, the critical timeframe for which males store sperm before fertilisation will depend on the trade-off between effects on sperm quantity versus quality. For example, when entire ejaculates are used to fertilise eggs, such as in IVF (e.g. Van Voorhis et al, 2001), the number of high-quality sperm would need to be maximised. Here, a meta-analysis including both sperm quality and quantity changes during sperm storage is needed predict the precise timing of sperm storage that maximizes the numbers of high-quality sperm. On the other hand, when single sperm are selected and directly used for fertilisation such as in ICSI, only sperm quality needs to be optimised (Devroey and Steirtghem, 2004). In such cases, shorter durations of sperm storage would be recommended.

Our study only included data on in-vivo sperm storage. However, in-vitro sperm storage via cryopreservation or sperm extenders is commonly used for artificial insemination and in sperm banks (Contreras et al, 2020; Sharafi et al, 2022). Future meta-analysis comparing our results on in-vivo (natural) rates of sperm senescence against in-vitro rates, could test whether artificial techniques are more efficient at preserving sperm than evolved biological mechanisms. Such comparisons might have crucial implications for improving sperm preservation techniques and inventing new biomimetic technologies to improve sperm storage.

## Conclusions

Our study documents the magnitude of how sperm storage, despite its potential evolutionary benefits, leads to declines in sperm quality and certain post-mating and post-fertilization outcomes in human and non-human animals. We further identify specific biological variables such as the sex of storage and type of trait, as well as methodological factors such as the experimental design and duration of storage, that modulate observed outcomes. We discuss how the specific physiology of sperm production, sperm recycling, and storage organs might mediate the magnitude of post-meiotic sperm senescence. Our results provide insights into the mechanisms that cause sperm deterioration during storage and suggestions for adaptive responses that animals might have evolved to ameliorate this. Importantly, we highlight the need to understand how natural selection might act on sperm in age-structured ejaculates. By collating data from both zoological and biomedical fields, combining evolutionary and physiological theories, and testing the influence of multiple variables, we provide generalisable and useful insights into the biology of sperm storage. Given the ubiquity of sperm storage across animals, we hope that our study generates greater interest in this overlooked topic, leading to further investigation of the mechanisms which facilitate long-term sperm storage and the costs or benefits of sperm storage.

## Author contributions

Conceptualisation- KS, RD, IS, RVT

Methodology- KS, RD, RVT, SN

Paper screening- KS, RD, KR

Data curation- KS, RD

Data validation- RD, KS

Formal analysis- KS, RD, RVT

Visualization- RD, RVT, KS

Writing (original draft)- KS, RD

Writing (revision)- RVT, IS, KR, SN

Funding acquisition- RD, IS, KS, RVT

## Acknowledgements and funding

We thank Tom Pizzari and Simone Immler for helpful discussions and comments. We are grateful to Hiroki Takeuchi, Tara DeLecce, and Yuchuan Zhou, for sending us missing data. KS was supported by an SSE Rosemary Grant award and ASN graduate student award. RD was supported by a Daphne Jackson Fellowship (NERC). IS was supported by a BBSRC Fellowship (BB/T008881/1) and a Royal Society Dorothy Hodgkin Fellowship (DHF\R1\211084). RVT was supported by a BBSRC standard grant awarded to Tommaso Pizzari (BB/V001256/1).

## Ethics statement

Our study did not use any live organisms or tissues, and obtained data on humans from publicly available sources that did not include any personal information.

## Data availability

All data associated with our study, and the code used to analyse this data, is available at OSF https://osf.io/jxv7z/files/osfstorage with the DOI: 10.17605/OSF.IO/JXV7Z.

## Supplementary information

Appendix 1-9, Supplementary Figures S1-S10 are provided in the Supplementary information

