## Supplementary information for "Sperm storage reduces sperm and embryo quality in animals"

**Appendix 1: Search string and PRISMA diagram**

We created a word cloud from the abstracts and titles of 20 relevant papers on post-meiotic sperm ageing and used this word cloud to generate an appropriate search string to obtain studies addressing the topic of our meta-analysis. Our search strings were:

SCOPUS (21/9/22): *( TITLE-ABS-KEY (* meiotic* *)  OR  TITLE-ABS-KEY (* storage *)  OR  TITLE-ABS-KEY (* senescence *)  OR  TITLE-ABS-KEY (* stored *)  AND  TITLE-ABS-KEY (* sperm *)  OR  TITLE-ABS-KEY (* ejaculate* *)  AND NOT  TITLE-ABS-KEY (* vitro *)  AND NOT  TITLE-ABS-KEY (* freez* *)  AND NOT  TITLE-ABS-KEY (* cold *)  AND NOT  TITLE-ABS-KEY (* frozen *)  AND NOT  TITLE-ABS-KEY (* patient *)  AND NOT  TITLE-ABS-KEY (* men *) ).*

Web of science (12/10/22): *TI= ((post meiotic OR storage OR senescence OR stored) AND (sperm OR ejaculate*) NOT (vitro* OR in-vitro OR freeze OR extender OR extenders OR chilled OR cryo* OR cryopreserved OR cryopreservation OR spermatid OR spermatogenesis OR spermatogonia OR spermiogenesis OR patient OR patients OR prophase OR anaphase OR metaphase OR cold OR cool OR frozen OR men OR human OR refrigerate OR refrigerated OR refrigeration OR freezing)) OR AB= ((post meiotic OR storage OR senescence OR stored) AND (sperm OR ejaculate*) NOT (vitro* OR in-vitro OR freeze OR extender OR extenders OR chilled OR cryo* OR cryopreserved OR cryopreservation OR spermatid OR spermatogenesis OR spermatogonia OR spermiogenesis OR patient OR patients OR prophase OR anaphase OR metaphase OR cold OR cool OR frozen OR men OR human OR refrigerate OR refrigerated OR refrigeration OR freezing)).*

In humans, we similarly used a word cloud generated from studies relevant to sperm storage in men, to create a search string to obtain relevant studies for screening. Our search strings were:

SCOPUS (26/01/2024): *TITLE-ABS-KEY ( abstinence AND ( sperm OR semen ) ) AND ( LIMIT-TO ( LANGUAGE , "English" ) )*

Web of science (26/01/2024): ***((TI=(abstinence)) AND (TI=(sperm) OR TI=(semen))) OR ((AB=(abstinence)) AND (AB=(sperm) OR AB=(semen)))***

After applying our inclusion and exclusion criteria, and screening abstracts and full-texts, we obtained 115 relevant human studies as well as 56 animal studies encompassing 30 species from which we collected data for our meta-analysis (Figure S1).

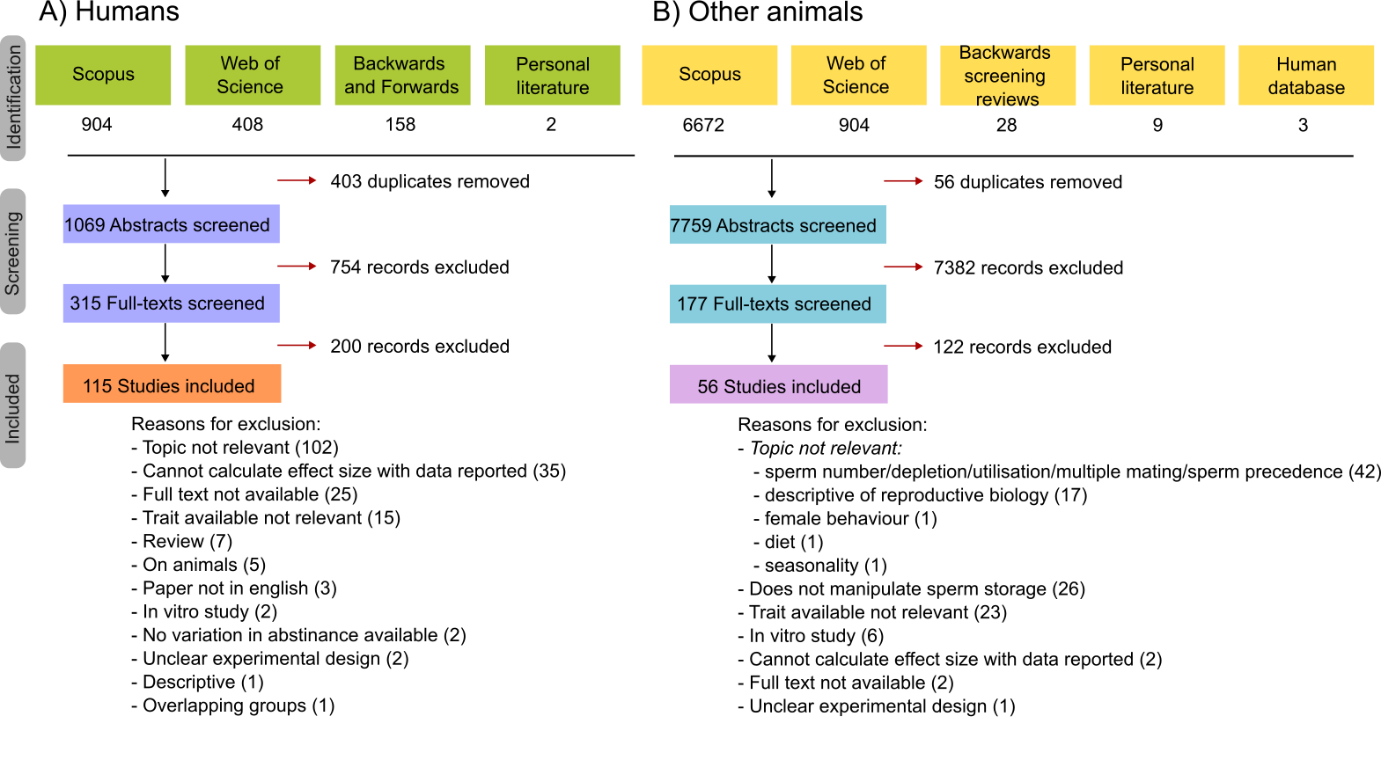

Figure S1: PRISMA diagram of the screening of (A) human and (B) animal studies, representing the source of studies, the number of studies excluded at each stage and the reasons for doing so. Personal literaature represents other sources, such as existing literature in the analysts paper collection, papers obtained from Google Scholar, or PhD theses.

**Appendix 2: Backwards and forwards search**

List of papers used for backwards and forwards screening to obtain additional human studies:

- Ayad, B.M., Van der Horst, G. and Du Plessis, S.S., 2017. Revisiting the relationship between the ejaculatory abstinence period and semen characteristics. *International journal of fertility & sterility*, *11*(4), p.238.
- Barbagallo, F., Cannarella, R., Crafa, A., La Vignera, S., Condorelli, R.A., Manna, C. and Calogero, A.E., 2023. The impact of a very short abstinence period on assisted reproductive technique outcomes: A systematic review and meta-analysis. *Antioxidants*, *12*(3), p.752.
- Barbagallo, F., Cannarella, R., Crafa, A., Manna, C., La Vignera, S., Condorelli, R.A. and Calogero, A.E., 2022. The impact of a very short abstinence period on conventional sperm parameters and sperm DNA fragmentation: a systematic review and meta-analysis. *Journal of Clinical Medicine*, *11*(24), p.7303.
- Du, C., Li, Y., Yin, C., Luo, X. and Pan, X., 2024. Association of abstinence time with semen quality and fertility outcomes: a systematic review and dose–response meta‐analysis. *Andrology*.
- Hanson, B.M., Aston, K.I., Jenkins, T.G., Carrell, D.T. and Hotaling, J.M., 2018. The impact of ejaculatory abstinence on semen analysis parameters: a systematic review. *Journal of assisted reproduction and genetics*, *35*, pp.213-220.
- Sokol, P., Drakopoulos, P. and Polyzos, N.P., 2021. The effect of ejaculatory abstinence interval on sperm parameters and clinical outcome of ART. A systematic review of the literature. *Journal of clinical medicine*, *10*(15), p.3213.
- Sørensen, F., Melsen, L.M., Fedder, J. and Soltanizadeh, S., 2023. The influence of male ejaculatory abstinence time on pregnancy rate, live birth rate and DNA fragmentation: a systematic review. *Journal of Clinical Medicine*, *12*(6), p.2219.

List of papers used for backwards screening to obtain additional animal studies:

- Austin C R. 1985. Sperm maturation in the male and female genital tracts. Pages 121–155 in Biology of Fertilization, Volume 2: Biology of the Sperm, edited by C B Metz and A Monroy. Orlando (FL): Academic Press.
- Bishop D W. 1962. Sperm motility. Physiological Reviews 42:1–59.
- Dziuk P J. 1996. Factors that influence the proportion of offspring sired by a male following heterospermic insemination. Animal Reproduction Science 43:65–88
- Lanman J T. 1968. Delays during reproduction and their effects on the embryo and fetus. New England Journal of Medicine 278:993–999.
- Orgebin-Crist M-C. 1973. Sperm age: effects on zygote development. Pages 85–95 in Proceedings of a Research Conference on Family Planning, edited by W A Uricchio. Washington (DC): Human Life Foundation.
- Pizzari T, Dean R, Pacey A, Moore H, Bonsall MB. The evolutionary ecology of pre- and post-meiotic sperm senescence. Trends Ecol Evol 23: 131-140
- Reinhardt K 2007. Evolutionary consequences of sperm cell aging. Quarterly Review of Biology 82: 375-393
- Reinhardt K, Köhler G, Schumacher J. 1999. Females of the grasshopper *Chorthippus parallelus* (Zett.) do not remate for fresh sperm. Proceedings of the Royal Society of London B 266:2003–2009.
- Salisbury G W, Hart R G. 1970. Gamete aging and its consequences. Biology of Reproduction (Supplement) 2:1–13.
- Siva-Jothy M T. 2000. The young sperm gambit. Ecology Letters 3:172–174.
- Tarín J J, Pérez-Albalá S, Cano A. 2000. Consequences on offspring of abnormal function in ageing gametes. Human Reproduction Update 6:532–549.

**Appendix 3: Collecting data on means and errors**

We collected data on means, SD/SE, and sample sizes of sperm-storing individuals in each storage group from: text in the results section, figures using WebPlotDigitizer (Rohtagi, 2014), the supplementary information or raw data provided with the paper, a data repository, or by directly contacting the authors of the paper, in that order. We converted standard errors to standard deviations (SD) using the formula: $SD=SE\cdot\sqrt{N}$. When medians and inter-quartile ranges were reported, we converted these to SD using Wan et al (2014).

**Appendix 4: Repeatability**

To ensure repeatability of data extraction for human studies, two analysts- KS and RVT, checked data obtained from >10% of studies (*i.e.* 15 studies). To do this, analyst 1 and 2 first independently extracted data and calculated effect sizes (Fisher’s z-transformed correlation coefficient (Zr); 57 effect sizes across 15 papers) of the data collected. Next, we constructed a linear mixed model, with Zr values of analyst 1 as the dependent variable, Zr values of analyst 2 as the predictor variable, and paper ID as a random effect. Then we tested for repeatability of Zr (effect size) and its variance (VZr) calculated by analyst 1 and 2 from collected data. This was: marginal R^2^ = 0.939 for Zr and 0.987 for VZr, indicating strong repeatability. Similarly, to ensure repeatability of data extraction for animal studies, the two analysts checked data obtained from 11 papers across 54 effect sizes. To do this, analyst 1 and 2 first independently extracted data and calculated effect sizes (Zr) and their variances (VZr). Next, we constructed a linear mixed model, with Zr values of analyst 1 as the dependent variable, Zr values of analyst 2 as the predictor variable, and paper ID as a random effect. Then a coefficient of determination of this model between the effect sizes obtained by analyst 1 and 2 was calculated (marginal R^2^ = 0.917 for Zr and 0.496 for VZr). The two analysts discussed the reasons for discrepancies in VZr values. These were caused by differences in how sample sizes were being recorded. Analysts agreed that sample sizes should be recorded as the number of individuals of the sperm-storing sex, and discrepant data was corrected accordingly.

**Appendix 5: Definitions and categorisations of traits**

| **Broad trait** | **Trait** | **Study-specific descriptions** |
| --- | --- | --- |
| Sperm  intra-cellular quality | DNA quality | - % sperm with fragmented DNA |
|  |  | - % DNA integrity- DNA fragmented grade 2 |
|  |  | - % DNA integrity- DNA fragmented grade 3 |
|  |  | - % DNA integrity- DNA fragmented grade 4 |
|  |  | - % total chromosomal abnormalities |
|  |  | - DNA integrity |
|  |  | - sperm chromatin structure damage-  based on fluorescence intensity |
|  | Sperm  oxidative  stress | - % superoxidase anion DHE positive |
|  |  | - 8-OHdG % sperm |
|  |  | - Fluorescence lifetimes |
|  |  | - ROS production rate % |
|  |  | - ROS rate controlled for volume of sperm |
|  |  | - Antioxidant capacity |
|  |  | - catalase O2- ANTIOXIDANT production |
|  |  | - lipid peroxidation (malondialdehyde level nM/hr) |
|  |  | - oxidative stress (level 3 and 4) |
|  |  | - oxidative stress adduct - amount of ROS (uM) |
|  |  | - pro-oxidants ROS production |
|  |  | - Reactive oxygen species % |
|  |  | - ROS (RLU/10^6 sperm) |
|  |  | - sod O2- ANTIOXIDANT production |
|  |  | - sperm O2- ROS production |
|  |  | - Sperm protein oxidation (nmol PC/mg protein) |
|  |  | - tbars O2- ROS production |
|  |  | - Total antioxidant capacity (mM) |
| Sperm  performance  Sperm  performance | Fertilisation  success | - % embryos fertilised after controlling for sperm number |
|  |  | - % fertilisation success while controlling for sperm number |
|  |  | - % implantation rate |
|  |  | - % pregnancy rate |
|  |  | - Fertilisation rate (ICSI)- % embryos fertilised  where one sperm inseminated in one egg |
|  |  | - % inseminated females fertilised (standardised sperm quantity used) |
|  |  | - % ova fertilised (standardised sperm quantity used) |
|  |  | - Proportion of fertilised eggs |
|  |  | - Proportion of fertilised eggs by 2nd male (fresh)  compared to first male (fresh or stored)- sperm defence |
|  |  | - Sperm ejection by female probability |
|  |  | - Number of offspring produced,  after controlling for sperm number |
|  |  | - Paternity share under sperm competition, after controlling for sperm number |
|  |  | - % acrosome reacted sperm |
|  |  | - Implantation rate % |
|  |  | - sperm acrosome reaction test |
|  | Sperm  motility | - % forward motility |
|  |  | - % immotile |
|  |  | - % immotile- grade a, d |
|  |  | - % motile |
|  |  | - % progressive |
|  |  | - Progressive motility on a subjective scale |
|  |  | - % rapid forward motility |
|  |  | - % rapid progressive |
|  |  | - % slow progressive motility |
|  |  | - % total motility |
|  | Sperm  velocity | - mean velocity |
|  |  | - mean velocity of motile sperm |
|  |  | - VAP, VCl, or VSL |
|  | Sperm  viability | - % acrosome integrity |
|  |  | - In-vitro longevity (time until movement stops) after in-vivo storage |
|  |  | - % membrane integrity |
|  |  | - % viable (live) sperm |
|  |  | - % vitality |
|  |  | - eosin-negrosin % viable |
|  |  | - Hypo-osmotic swelling test |
|  |  | - Membrane integrity |
|  |  | - propidium iodide test |
|  | Sperm  morphology | - % sperm with abnormal morphology |
|  |  | - % morphological defects sperm- grade 3 |
|  |  | - % sperm with normal morphology |
|  |  | - % normal acrosomes |
|  |  | - % normal head morphology sperm |
|  |  | - % normal morphology sperm |
|  |  | - % normal sperm morphology |
|  |  | - % normal tail morphology sperm |
|  |  | - % tail abnormalities |
|  |  | - % normal midpiece morphology sperm |
| Post- fertilisation | Embryo  quality | - % abortion rate |
|  |  | - % anueploid embryos |
|  |  | - % normal blastocyst development |
|  |  | - % embryos observed with cleavage |
|  |  | - % embryos with blastocyst development |
|  |  | - % euploid embryos (correct number of chromosomes) |
|  |  | - % grade A top quality embryos |
|  |  | - % high quality blastocysts |
|  |  | - % high quality day 3 blastocysts |
|  |  | - % high quality day 3 embryos |
|  |  | - % miscarriage rate |
|  |  | - % motile |
|  |  | - % top quality embryos on day 3 |
|  |  | - blastulation rate |
|  |  | - Early miscarriage rate %:  % miscarriages divided by total successful pregnancies |
|  |  | - embryo score |
|  |  | - % Live birth rate |
|  |  | - % fertilised eggs that hatched |
|  |  | - oocyte cleavage rate (no. of cleaved oocytes) |
|  |  | - oocyte maturation rate % |
|  | Post-hatching offspring  quality | - Offspring body size |
|  |  | - Offspring fecundity |
|  |  | - Offspring survival |
|  |  | - Sons' sperm quality and quantity |
|  |  | - % offspring surviving until X days |
|  |  | - Offspring paternity share |
|  |  | - Offspring fertility |
|  |  | - % embryos with chromosomal abnormalities |

**Appendix 6: absolute sperm storage durations- range, max, min, for humans and animals**

Minimum (X axis) and maximum duration (Y axis) for which sperm were stored in individuals within a study, plotted against each other to show the distribution of effect sizes for these durations, and the range of sampling (i.e. maximum – minimum duration of storage) for each study. Range of sampling was highly skewed for both humans (Figure S2), and animal studies (Figure S3). Therefore, these values were log_e_ transformed to allow for a more normal distribution of data, before being used for our analysis.

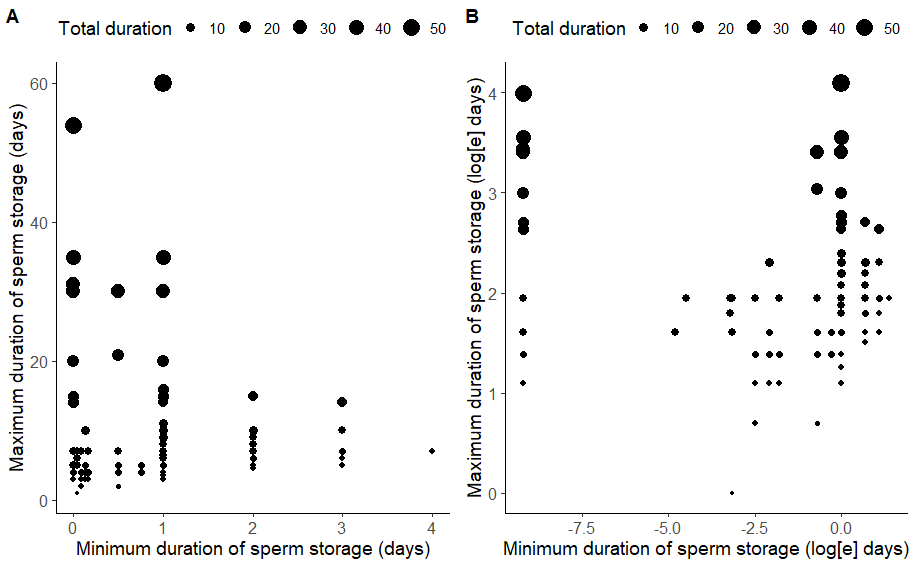

Figure S2: Minimum, maximum, and total duration of abstinence/sperm storage (A- days, B- log_e_ transformed days) sampled by a study in the human dataset. Size of points (effect size) represents the range (i.e. maximum – minimum duration) of storage duration sampled by a study. One study was excluded as this contained males with spinal cord injuries and had an unusually high sexual abstinence range (i.e. > 2000 days). 1 hour = -3.18 log_e_ days, 1 day = 0 log_e_ days, 10 days = 2.3 log_e_ days, 50 days = 3.9 log_e_ days.

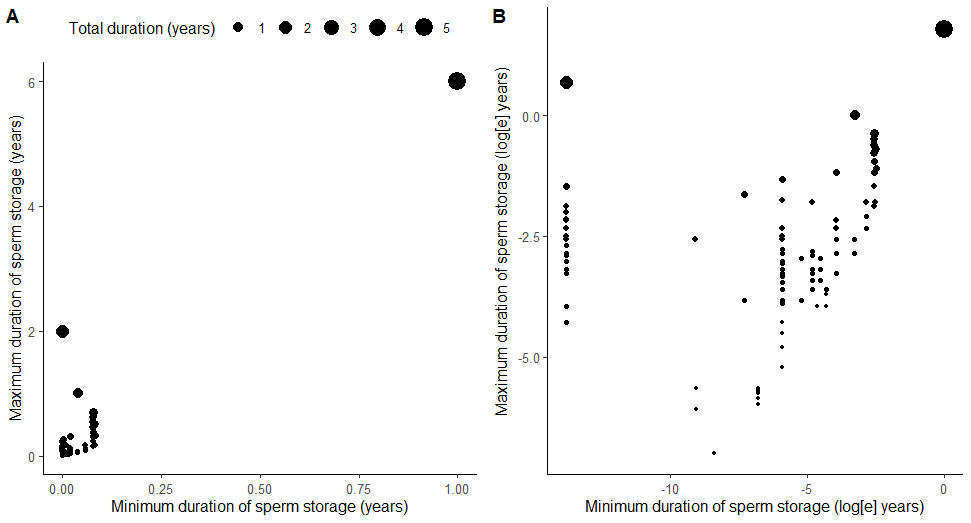

Figure S3: Minimum, maximum, and total duration of sperm storage (A- years, B- log_e_ transformed years) sampled by a study in the animal dataset. Size of points (effect size) represents the range (i.e. maximum – minimum duration) of storage duration sampled by a study. 1 hour = -9.079 log_e_ years, 1 day = -5.9 log_e_ years, 1 week = -3.96 log_e_ years, 1 month = -2.485 log_e_ years, 1 year = 0 log years, 5 years = 1.61 log_e_ years.

**Appendix 7: Zr calculation**

We first calculated r (correlation coefficient) using different methods and then used r to calculate Zr.

A. Two sperm storage groups (Nakagawa and Cuthill, 2007)

To calculate correlation coefficients from studies which reported comparisons between two sperm storage groups, we first calculated a standardized mean difference (SMD). SMD here, was calculated using the package *metafor* in R with the function *escalc.* SMD provides the strength of an effect by dividing the difference in means between two groups, by their pooled standard deviation. We used the following formula:

SMD = $\frac{\mu_{long} -\mu_{short}}{S pooled}$

$$S_{pooled}=\sqrt{\frac{\left( n_{long}-1 \right)S_{long}^{2} + \left( n_{short}-1 \right)S_{short}^{2}}{n_{long} + n_{short} - 2}}$$

where $\mu_{long}$ and $\mu_{short}$ are means of the trait values for long and short sperm storage groups respectively, and $S_{pooled}$ is the pooled standard deviation. These SMD values were then converted to a correlation coefficient (following Borenstein et al, 2009; Polanin and Snilstveit, 2016) using the formula:

$$r=\frac{ⅆ}{\sqrt{\frac{d^{2}+1}{\frac{N_{long}}{N_{short}+N_{long}}\cdot\left( 1-\frac{N_{long}}{N_{short}+N_{long}} \right)}}}$$

Where $N_{long}$and $N_{short}$are the unique sample sizes of sperm-storing individuals in the long and short storage groups.

B. Multiple (>2) sperm storage groups

To calculate effect sizes from studies that reported means and standard deviations from more than two sperm storage groups, we used a simulation (with 1000 iterations). This simulation calculated a correlation coefficient between storage duration groups and their means, while weighting the means by their standard deviations (following Johnson et al, 2015; Sanghvi et al, 2024). To test for consistency between effect size outcomes from the simulation and the outcomes from SMD, we additionally calculated correlation coefficients using a simulation for outcomes with only two-groups. There was strong agreement between r values obtained from these two methods (i.e. SMD and simulation) ($R^{2}$> 0.95).

C. Test statistics

In the absence of means and SD/SE, we collected data from test-statistics and used these to calculate point biserial correlation coefficients ($r_{pb}$) (following Borenstein et al, 2009; Koricheva et al, 2013; Polanin and Snilstveit, 2016). We then converted $r_{pb}$ to a biserial correlation coefficient ($r_{b})$.

1. For converting Beta (regression coefficient) to t (Peterson and Brown, 2005)

$$t=\frac{\beta}{SE}$$

Where SE is the standard error

2. For converting “t” from independent t-test into $r_{pb}$

$$r_{pb} =\frac{t}{\sqrt{(t^{2}}+DF)}$$

Where DF is the degrees of freedom

3. For converting F from ANOVA or ANCOVA with 1 degree of freedom to $r_{pb}$

$$r_{pb} =\sqrt{(\frac{F}{F+N-2})}$$

Where N is the sample size of unique number of sperm-storing individuals

4. For converting spearman’s rho to *r*

$$r =2*\sin((\pi* \rho)/6)$$

5. For converting Odd’s ratio to correlation coefficient

$$r=\frac{\ln\left( OR \right)\cdot\left( \frac{\sqrt{3}}{\pi} \right)}{\sqrt{\left( \ln\left( OR \right) \cdot\frac{\sqrt{3}}{\pi} \right)^{2}}+4}$$

6. For converting R squared and adjusted R squared values to *r*

$$r= \sqrt{R^{2}}$$

7. For converting $\boldsymbol{\chi}^{2}$ values from Chisq. test with one degree of freedom, to $r_{pb}$

$$r_{pb} = \sqrt{\frac{\boldsymbol{\chi}^{2}}{N}}$$

8. For converting Pearson’s correlation coefficient (r) to *r*

$$r=r_{Pearson's}$$

9. Point biserial to biserial (following Jacobs and Viechtbauer, 2016; McCartney et al, 2024)

$$r=r_{b}=r_{pb}\cdot\left( \frac{\sqrt{\frac{N_{long}}{N_{short}+N_{long}}\cdot\frac{N_{short}}{N_{short}+N_{long}}}}{\phi^{-1}\cdot\left( \phi\cdot\left( \frac{N_{long}}{N_{short}+N_{long}} \right) \right)} \right)$$

D. When means of groups were reported as % without SD or SE

When only the means of two storage groups without SD or SE were reported, and the means were reported as a % (e.g. % of motile sperm, % of viable sperm, % of surviving embryos), we used the following formula (Hasselblad and Hedges, 1995):

$$r=\frac{\frac{logit\left( \frac{\mu_{long}}{100} \right)-logit\left( \frac{\mu_{short}}{100} \right)}{\frac{\pi}{\sqrt{3}}}}{\sqrt{{\frac{logit\left( \frac{\mu_{long}}{100} \right)-logit\left( \frac{\mu_{short}}{100} \right)}{\frac{\pi}{\sqrt{3}}}}^{2}+\frac{1}{\frac{N_{long}}{N_{short}+N_{long}}\cdot\left( 1-\frac{N_{long}}{N_{short}+N_{long}} \right)}}}$$

In such scenarios, when there were multiple (i.e. >2) storage groups, we conducted pairwise comparisons between the shortest sperm storage groups against each of the other groups using the above formula.

E. Correlation coefficients (r) to Fisher’s z transformed correlation coefficient (Zr):

$$Z_{r}=0.5\cdot(log \left( 1+r \right)-\log\left( 1-r \right))$$

**Appendix 8: Effective sample size**

We calculated effective sample sizes $N_{e}$ following Rutkowska et al (2014) as:

$$N_{e}=\frac{N_{total}}{1+\left( \frac{N_{total}}{N_{unique}} -1 \right) \cdot0.8}$$

Where $N_{total}$ was the total number of sperm-storing individuals (male or female) sampled across all treatments, including individuals that were repeatedly sampled. $N_{unique}$was the unique number of individuals sampled. For longitudinal and semi-longitudinal studies, $N_{total}>N_{unique}$, for cross-sectional studies $N_{total}=N_{unique}$.

**Appendix 9: list of included studies**

Humans:

| Paper ID | Title | DOI or web link |
| --- | --- | --- |
| H1 | Influence of the abstinence period on human sperm quality: analysis of 2,458 semen samples | 10.5935/1518-0557.20170052 |
| H10 | Can a short term of repeated ejaculations affect seminal parameters? | https://www.ncbi.nlm.nih.gov/pmc/articles/PMC4947206/ |
| H100 | Intelligence and semen quality are positively correlated | 10.1016/j.intell.2008.11.001 |
| H101 | No evidence for a trade-off between competitive traits and ejaculate quality in humans | https://doi.org/10.1177/1474704920942557 |
| H102 | Nonspermatozoal cells in human sperm: A study of 1243 subfertile and 253 fertile men | 10.3109/01485018409161176 |
| H103 | Sperm donor lifestyle survey: modifiable risk factors for potential sperm donors | 10.1007/s10815-021-02322-x |
| H104 | Trends in sperm quality by computer-assisted sperm analysis of 49,189 men during 2015-2021 in a fertility center from China | 10.3389/fendo.2023.1194455 |
| H105 | Influence of abstinence period on clinical outcomes in fresh embryo transfer after intracytoplasmic sperm injection. | 10.1016/j.fertnstert.2015.07.914 |
| H106 | Paternal contribution to embryonic competence | 10.5173/ceju.2019.1900 |
| H107 | The effect of ejaculatory frequency on semen characteristics of normozoospermic and oligozoospermic men from an infertile population | 10.1093/oxfordjournals.humrep.a137877 |
| H108 | Mate retention behavior and ejaculate quality in humans | 10.1007/s10508-021-01992-z |
| H109 | Age, sexual abstinence duration, sperm morphology, and motility are predictors of sperm DNA fragmentation | https://doi.org/10.1002/rmb2.12585 |
| H11 | Changes in seminal parameters of ejaculates after repeated ejaculation | https://doi.org/10.1111/j.1439-0272.1988.tb02363.x |
| H110 | Association of lifestyle and occupational exposure factors with human semen quality: a cross-sectional study of 1060 participants | https://doi.org/10.1080/19396368.2024.2357348 |
| H111 | No Evidence for a Relationship between Intelligence and Ejaculate Quality | 10.1177/1474704920960450 |
| H112 | Preliminary prediction of semen quality based on modifiable lifestyle factors by using the XGBoost algorithm | https://doi.org/10.3389/fmed.2022.811890 |
| H113 | Lifestyle and demographic factors associated with human semen quality and sperm function | https://doi.org/10.1080/19396368.2018.1491074 |
| H114 | Decline in Semen Quality among Fertile Men in Paris during the Past 20 Years | 10.1056/NEJM199502023320501 |
| H115 | Early and late paternal contribution to cell division of embryos in a time-lapse imaging incubation system | https://doi.org/10.1111/and.14211 |
| H12 | Characterization of the sperm proteome and reproductive outcomes with in vitro fertilization after a reduction in Male ejaculatory abstinence period | 10.1074/mcp.RA117.000541 |
| H13 | Comparison of sperm parameters, in vitro fertilization results, and subsequent pregnancy rates using sequential ejaculates, collected two hours apart, from oligoasthenozoospermic men | 10.1016/S0015-0282(16)57920-5 |
| H14 | Correlation between sperm motility and sperm chromatin structure assay parameters | 10.1016/S0015-0282(03)02212-X |
| H15 | Determination of active, non-zymogen acrosin, proacrosin and total acrosin in different andrological patients | 10.1007/BF00375729 |
| H16 | Does duration of abstinence affect the live-birth rate after assisted reproductive technology? A retrospective analysis of 1,030 cycles | 10.1016/j.fertnstert.2017.08.034 |
| H17 | Duration of sexual abstinence: epididymal and accessory sex gland secretions and their relationship to sperm motility | https://doi.org/10.1093/humrep/deh586 |
| H18 | Effect of abstinence duration on sperm sex ratio | https://he02.tci-thaijo.org/index.php/sirirajmedj/article/view/55267 |
| H19 | Effect of abstinence on sperm acrosin, hypoosmotic swelling, and other semen variables | https://doi.org/10.1016/S0015-0282(16)55330-8 |
| H2 | A comparative evaluation of semen parameters in pre- and post-Hurricane Katrina human population | 10.4103/1008-682X.143738 |
| H20 | Effect of abstinence on sperm motility in normal men | https://doi.org/10.1016/0002-9378(88)90038-5 |
| H21 | Effect of abstinence time on semen parameters among male patients referring to urology clinic | 10.9734/JPRI/2019/v31i330300 |
| H22 | Effect of age and abstinence on semen quality: A retrospective study in a teaching hospital | 10.1016/S2305-0500(14)60017-10 |
| H23 | Effect of duration of abstinence on maturity of human spermatozoa nucleus | https://doi.org/10.3109/01485018608986954 |
| H24 | Effect of ejaculatory abstinence period on fertilization and clinical outcomes in ICSI cycles: a retrospective analysis | https://doi.org/10.1016/j.rbmo.2023.103401 |
| H25 | Effect of ejaculatory abstinence period on sperm DNA fragmentation and pregnancy outcome of intrauterine insemination cycles: A prospective randomized study | https://doi.org/10.1007/s00404-020-05783-0 |
| H26 | Effect of time interval between ejaculations on semen parameters | 10.3109/01485019108987658 |
| H27 | Effects of a short abstinence period on sperm quality in oligozoospermic men | https://doi.org/10.5653/cerm.2023.06100 |
| H28 | Effects of multiple ejaculations after extended periods of sexual abstinence on total, motile and normal sperm numbers, as well as accessory gland secretions, from healthy normal and oligozoospermic men | 10.1093/oxfordjournals.humrep.a138236 |
| H29 | Effects of varying the abstinence period in the same individuals on sperm quality | 10.3109/01485019108987644 |
| H3 | Abstinence time and its impact on basic and advanced semen parameters | 10.1016/j.urology.2016.03.059 |
| H30 | Ejaculatory abstinence affects the sperm quality in normozoospermic men-how does the seminal bacteriome respond? | 10.3390/ijms24043503 |
| H31 | Evaluation of semen quality in 1808 university students, from Wuhan, Central China | 10.4103/1008-682X.135984 |
| H32 | Evidence for obtaining a second successive semen sample for intrauterine insemination in selected patients: results from 32 consecutive cases | 10.5653/cerm.2016.43.2.102 |
| H33 | Exposures that may affect sperm DNA integrity: Two decades of follow-up in a pregnancy cohort | 10.1016/j.reprotox.2011.12.013 |
| H34 | How 1 h of abstinence improves sperm quality and increases embryo euploidy rate after PGT-A: a study on 106 sibling biopsied blastocysts | 10.1007/s10815-019-01533-7 |
| H35 | How well do semen analysis parameters correlate with sperm DNA fragmentation? a retrospective study from 2567 semen samples analyzed by the halosperm test | https://doi.org/10.3390/jpm13030518 |
| H36 | Impacts of abstinence time on semen parameters in a large population-based cohort of subfertile men | https://doi.org/10.1016/j.urology.2017.06.045 |
| H37 | Impact of ejaculatory abstinence period and semen characteristic on the reproductive outcomes after intracytoplasmic sperm injection | 10.5935/1518-0557.20210124 |
| H38 | Does a very short length of abstinence improve assisted reproductive technique outcomes in infertile patients with severe oligo-asthenozoospermia? | https://doi.org/10.3390/jcm10194399 |
| H39 | High percentage of abnormal semen parameters in a prevasectomy population | https://doi.org/10.1016/j.fertnstert.2005.09.032 |
| H4 | Analysis of human sperm DNA fragmentation index (DFI) related factors: a report of 1010 subfertile men in China | https://doi.org/10.1186/s12958-018-0345-y |
| H40 | Impact of geographical and seasonal temperature on sperm parameters in Indian men who were partners in subfertile couples – A retrospective analysis | 10.4103/2305-0500.350153 |
| H41 | Impact of the sexual abstinence period on the production of seminal reactive oxygen species in patients undergoing intrauterine insemination: A randomized trial | https://doi.org/10.1111/jog.14308 |
| H42 | Improved sperm DNA fragmentation levels in infertile men following very short abstinence of 3–4 hours | https://dx.doi.org/10.21037/tau-23-216 |
| H43 | Improved sperm kinematics in semen samples collected after 2 h versus 4–7 days of ejaculation abstinence | https://doi.org/10.1093/humrep/dex101 |
| H44 | Improved sperm motility after 4 h of ejaculatory abstinence: role of accessory sex gland secretions | https://doi.org/10.1071/RD18135 |
| H45 | Short-interval second ejaculation improves sperm quality, blastocyst formation in oligoasthenozoospermic males in ICSI cycles: a time-lapse sibling oocytes study | 10.3389/fendo.2023.1250663 |
| H46 | Increased count, motility, and total motile sperm cells collected across three consecutive ejaculations within 24 h of oocyte retrieval: implications for management of men presenting with low numbers of motile sperm for assisted reproduction | https://doi.org/10.1007/s10815-015-0509-z |
| H47 | Increased pregnancy after reduced male abstinence | https://doi.org/10.3109/19396368.2013.790919 |
| H48 | Increasing paternal age and ejaculatory abstinence length negatively influence the intracytoplasmic sperm injection outcomes from egg-sharing donation cycles | https://doi.org/10.1111/andr.12737 |
| H49 | Influence of abstinence and ejaculation-to-analysis delay on semen analysis parameters of suspected infertile men | https://doi.org/10.3109/01485018208990205 |
| H5 | Application of leukocyte subsets and sperm dna fragment rate in infertile men with asymptomatic infection of genital tract | 10.21037/apm-19-597 |
| H50 | Influence of ejaculation frequency on seminal parameters | https://doi.org/10.1186/s12958-015-0045-9 |
| H51 | Influence of ejaculatory abstinence on seminal total antioxidant capacity and sperm membrane lipid peroxidation | https://doi.org/10.1016/j.fertnstert.2014.05.039 |
| H52 | Influence of ejaculatory abstinence period on semen quality of 5165 normozoospermic and oligozoospermic Nigerian men: A retrospective study | 10.1002/hsr2.722 |
| H53 | One abstinence day decreases sperm DNA fragmentation in 90 % of selected patients | https://doi.org/10.1007/s10815-013-0089-8 |
| H54 | One day is better than four days of ejaculatory abstinence for sperm function | 10.1530/RAF-20-0018 |
| H55 | Optimum abstinence time for cryopreservation of semen in cancer patients | https://doi.org/10.1016/S0022-5347(01)67235-5 |
| H56 | Relationship between sexual abstinence duration and the acrosome index in teratozoospermic semen: analysis of 1800 semen samples | https://doi.org/10.1111/j.1439-0272.2006.00715.x |
| H57 | Retrospective analysis of the first collection versus the second collection in severe oligo-asthenoteratozoospermia cases in self-intracytoplasmic sperm injection patients | 10.4103/jhrs.jhrs_46_22 |
| H58 | Revisiting the impact of ejaculatory abstinence on semen quality and intracytoplasmic sperm injection outcomes | https://doi.org/10.1111/andr.12572 |
| H59 | Role of sequential semen samples in infertile men candidates for assisted reproduction: A prospective study | https://doi.org/10.1016/j.afju.2018.09.001 |
| H6 | Assessment of semen characteristics in consecutive ejaculates with short abstinence using SQA-V sperm quality analyzer. | https://doi.org/10.1016/j.fertnstert.2021.07.936 |
| H60 | Semen analysis in fertile patients undergoing vasectomy: reference values and variations according to age, length of sexual abstinence, seasonality, smoking habits and caffeine intake | 10.1590/s1516-31802005000400002 |
| H61 | Semen characteristics in consecutive ejaculates with short abstinence in subfertile males | https://doi.org/10.1016/j.rbmo.2015.11.021 |
| H62 | Seminal plasma metabolomics profiles following long (4–7 days) and short (2 h) sexual abstinence periods | https://doi.org/10.1016/j.ejogrb.2021.07.024 |
| H63 | Short abstinence may have paradoxical effects on sperms with different level of DNA integrity: a prospective study | 10.22037/uj.v18i.6515 |
| H64 | Short abstinence: A potential strategy for the improvement of sperm quality | https://doi.org/10.1016/j.mefs.2017.07.005 |
| H65 | Short ejaculatory abstinence in normozoospermic men is associated with higher clinical pregnancy rates in sub-fertile couples undergoing intra-cytoplasmic sperm injection in assisted reproductive technology: a retrospective analysis of 1691 cycles | 10.4103/jhrs.jhrs_235_20 |
| H66 | Shorter abstinence decreases sperm deoxyribonucleic acid fragmentation in ejaculate | https://doi.org/10.1016/j.fertnstert.2011.08.027 |
| H67 | Improvement of sperm motility by short-interval sequential ejaculation in oligoasthenozoospermic patients | https://api.semanticscholar.org/CorpusID:56346002 |
| H68 | Sperm characteristics in normal and abnormal ejaculates are differently influenced by the length of ejaculatory abstinence | https://doi.org/10.1111/andr.13222 |
| H69 | Sperm chromatin immaturity observed in short abstinence ejaculates affects DNA integrity and longevity in vitro | https://doi.org/10.1371/journal.pone.0152942 |
| H7 | Assessment of semen parameters in consecutive ejaculates with short abstinence period in oligospermic males | 10.5935/1518-0557.20210073 |
| H70 | Sperm parameters before and after swim-up of a second ejaculate after a short period of abstinence | https://doi.org/10.3390/jcm9041029 |
| H71 | Sperm parameters of the infertile patients in relation to socio demographic factors | [10.31838/ecb/2022.11.04.012](http://dx.doi.org/10.31838/ecb/2022.11.04.012) |
| H72 | Successful cryptozoospermia management with multiple semen specimen collection | https://doi.org/10.1016/j.fertnstert.2023.07.019 |
| H73 | The applicability of the flow cytometric sperm chromatin structure assay in epidemiological studies | 10.1093/humrep/13.9.2495 |
| H74 | The combination matters - distinct impact of lifestyle factors on sperm quality: a study on semen analysis of 1683 patients according to MSOME criteria | 10.1186/1477-7827-10-115 |
| H75 | The effect of age and abstinence time on semen quality a retrospective study | 10.4103/aja202165 |
| H76 | The effect of sperm DNA fragmentation index on assisted reproductive technology outcomes and its relationship with semen parameters and lifestyle | 10.21037/tau.2019.06.22 |
| H77 | The effects of short abstinence time on sperm motility, morphology and DNA mdamage | [10.4172/2167-0250.1000107](http://dx.doi.org/10.4172/2167-0250.1000107) |
| H78 | The impact of testicular and accessory sex gland function on sperm chromatin integrity as assessed by the sperm chromatin structure assay (SCSA) | https://doi.org/10.1093/humrep/17.12.3162 |
| H79 | The incidence and etiology of sperm DNA fragmentation in the ejaculates of males with spinal cord injuries | https://doi.org/10.1038/s41393-020-0426-6 |
| H8 | Association between the interval of sexual abstinence and sperm DNA fragmentation in patients with leukocytospermia | https://www.researchgate.net/publication/283969710 |
| H80 | The Variability of Semen Parameters With Sexual Abstinence Using Mail-in Sperm Testing Is Similar to That Seen With Traditional In-Office Semen Analysis | https://doi.org/10.1177/15579883231197910 |
| H81 | Three hour abstinence as a treatment for high sperm DNA fragmentation: a prospective cohort study | https://doi.org/10.1007/s10815-020-01999-w |
| H82 | Two subsequent seminal productions: A good strategy to treat very severe oligoasthenoteratozoospermic infertile couples | https://doi.org/10.1111/andr.13020 |
| H83 | Variability in the human-hamster in vitro assay for fertility evaluation | https://doi.org/10.1016/S0015-0282(16)46820-2 |
| H84 | Variation of semen measures within normal men | https://doi.org/10.1016/S0015-0282(16)48866-7 |
| H85 | Within-subject variability and the importance of abstinence period for sperm count, semen volume and pre-freeze and post-thaw motility | 10.1111/j.1439-0272.1981.tb00085.x |
| H86 | Within-subject variation of seminal parameters in men with infertile marriages | https://doi.org/10.1111/j.1365-2605.2006.00727.x |
| H87 | Relationship between the duration of sexual abstinence and semen quality: analysis of 9,489 semen samples | https://doi.org/10.1016/j.fertnstert.2004.12.045 |
| H88 | Analysis of semen parameters during 2 weeks of daily ejaculation: a first in humans study | 10.21037/tau.2016.08.20 |
| H89 | Antioxidant treatment of patients with asthenozoospermia or moderate oligoasthenozoospermia with high-dose vitamin C and vitamin E: a randomized, placebo-controlled, double-blind study | https://doi.org/10.1093/humrep/14.4.1028 |
| H9 | Basic phenotyping of male fertility from 2019 to 2020 at the human sperm bank of Fudan University | https://doi.org/10.1007/s43657-022-00047-0 |
| H90 | Comparison between length of sexual abstinence and semen parameters in patients of an assisted reproduction center and a hospital from porto alegre | 10.5935/1518-0557.2010.13.2.06 |
| H91 | Relationship between sexual abstinence of men and chromosomally abnormal spermatozoa | https://doi.org/10.1530/jrf.0.0910065 |
| H92 | Correlates of human sperm motility assessed by laser Doppler spectroscopy | 10.1111/j.1365-2605.1989.tb01280.x |
| H93 | Cross-sectional study of semen parameters in a large group of normal Chinese men | https://doi.org/10.1111/j.1365-2605.1985.tb00840.x |
| H94 | Effect of daily ejaculation on semen quality and calcium and magnesium in semen | https://doi.org/10.1016/j.androl.2013.03.001 |
| H95 | Effect of the sexual abstinence period recommended by the World Health Organization on clinical outcomes of fresh embryo transfer cycles with normal ovarian response after intracytoplasmic sperm injection | https://doi.org/10.1111/and.12964 |
| H96 | Ejaculatory abstinence in semen analysis: does it make any sense? | 10.1177/2633494120906882 |
| H97 | Factors affecting the variability of semen analysis results in infertile men | 10.1111/j.1365-2605.1981.tb00743.x |
| H98 | How often should infertile men have intercourse to achieve conception? | 10.1016/S0015-0282(16)56893-9 |
| H99 | Influence of the abstinence period on human sperm quality | 10.1016/j.fertnstert.2004.03.014 |

Animals:

| Paper ID | Title | DOI or weblink |
| --- | --- | --- |
| 1 | Male sperm storage compromises sperm motility in guppies | 10.1098/rsbl.2014.0681 |
| 2 | Sperm storage by males causes changes in sperm phenotype and influences the reproductive fitness of males and their sons | 10.1002/evl3.2 |
| 3 | Influence of female age, sperm senescence and multiple mating on sperm viability in female *Drosophila melanogaster* | 10.1016/j.jinsphys.2011.02.017 |
| 4 | The relationship of in vivo sperm storage interval to fertility and embryonic survival in the chicken | 10.1093/biolreprod/5.3.252 |
| 5 | Reduced metabolic rate and oxygen radicals production in stored insect sperm | https://doi.org/10.1098/rspb.2011.2422 |
| 6 | Male sperm storage impairs sperm quality in the zebrafish | https://doi.org/10.1038/s41598-021-94976-x |
| 7 | Females become infertile as the stored sperm's oxygen radicals increase | https://doi.org/10.1038/srep02888 |
| 8 | Sexual rest and post-meiotic sperm ageing in house mice | https://doi.org/10.1111/jeb.12661 |
| 9 | Sexual selection and ageing: Interplay between pre- and post-copulatory traits senescence in the guppy | https://doi.org/10.1098/rspb.2018.2873 |
| 10 | The effects of male age, sperm age and mating history on ejaculate senescence | ttps://doi.org/10.1111/1365-2435.13305 |
| 11 | The evolutionary consequences of sperm senescence in *Drosophila melanogaster* | http://hdl.handle.net/1974/8657 |
| 31 | Chromosome abnormalities in rabbit blastocysts resulting from spermatozoa aged in the male tract | https://doi.org/10.1016/S0015-0282(16)39556-5 |
| 32 | The effect of resting roosters from ejaculation on the quality of spermatozoa in semen | https://doi.org/10.1530/jrf.0.0110489 |
| 34 | Characteristics of the spermathecal contents of old and young honeybee queens | 10.1016/j.jinsphys.2008.10.010 |
| 35 | Functional association between female sperm storage organs and male sperm removal organs in calopterygid damselflies | 10.1111/j.1479-8298.2005.00123.x |
| 36 | An advantage for young sperm in the house cricket *Acheta domesticus* | 10.1086/430010 |
| 37 | Effect of sexual rest and subsequent regular collection on acrosome characteristics of bull spermatozoa | 10.2527/jas1970.31167x |
| 38 | Sperm aging in the male and cytogenetic anomalies. An animal model | https://doi.org/10.1007/BF00295606 |
| 39 | Sperm age, sex ratio, and hyperhaploidy frequency in mice | https://doi.org/10.1159/000133370 |
| 40 | Maturation and aging changes in rabbit spermatozoa isolated by ligatures at different levels of the epididymis | https://doi.org/10.1016/S0015-0282(16)37039-X |
| 44 | Effect of Stale Sperm on Fertility and Hatchability of Chicken Eggs | https://doi.org/10.3382/ps.0220218 |
| 45 | *Drosophila melanogaster* sperm turn more oxidative in the female | https://doi.org/10.1242/jeb.247775 |
| 46 | Factors affecting sperm quality before and after mating of calopterygid damselflies | https://doi.org/10.1371/journal.pone.0009904 |
| 47 | Haploid selection within a single ejaculate increases offspring fitness | https://doi.org/10.1073/pnas.1705601114 |
| 49 | Fertilization in the domestic fowl | https://doi.org/10.3382/ps.0080237 |
| 50 | A revised artificial insemination schedule for broiler breeder hens | <https://doi.org/10.3382/ps.0550725> |
| 51 | A study of the function of the epididymis: iii. functional changes undergone by spermatozoa during their passage through the epididymis and vas deferens in the guinea-pig | https://doi.org/10.1242/jeb.8.2.151 |
| 52 | The vitality of the spermatozoa in the male and female reproductive tracts | https://doi.org/10.1242/jeb.4.2.155 |
| 53 | No evidence for paternal age effects on sons or daughters, when accounting for paternal sperm storage | https://doi.org/10.1101/2024.09.19.613916 |
| 55 | Sperm competition and the function of masturbation in Japanese macaques (*Macaca fuscata*) | 10.5282/EDOC.105 https://edoc.ub.uni-muenchen.de/105/ |
| A12 | Viability of ram spermatozoa in relation to the abstinence period and successive ejaculations | 10.1111/j.1365-2605.1996.tb00477.x |
| A13 | Role of seminal MDA, ROS, and antioxidants in cryopreservation and their kinetics under the influence of ejaculatory abstinence in bovine semen | 10.1016/j.cryobiol.2020.11.002 |
| A14 | Parental age, gametic age, and inbreeding interact to modulate offspring viability in *Drosophila melanogaster* | 10.1111/evo.12131 |
| A15 | Sperm transfer and storage in relation to sperm competition in *Callosobruchus maculatus* | 10.1007/BF00171502 |
| A16 | Why do male *Callosobruchus maculatus* beetles inseminate so many sperm? | 10.1007/BF00175725 |
| A17 | The viability of hamster spermatozoa stored in the isthmus of the oviduct: The importance of sperm-epithelium contact for sperm survival | 10.1095/biolreprod42.3.450 |
| A22 | Extreme fertilization bias towards freshly inseminated sperm in a species exhibiting prolonged female sperm storage | 10.1098/rsos.172195 |
| A23 | Artificial insemination in Houbara bustards (*Chlamydotis undulata*): Influence of the number of spermatozoa and insemination frequency on fertility and ability to hatch | <https://doi.org/10.1530/jrf.0.1000093> |
| A24 | Copulatory behaviour increases sperm viability in female spiders | 10.1093/BIOLINNEAN/BLAA130 |
| A25 | Early events in seminal fluid and sperm storage in the female blue crab *Callinectes sapidus* Rathbun: Effects of male mating history, male size, and season | 10.1016/j.jembe.2005.01.001 |
| A26 | Function of multiple sperm storage organs in female Mediterranean fruit flies (*Ceratitis capitata*, Diptera: Tephritidae) | 10.1016/j.jinsphys.2004.11.007 |
| A27 | Decline in functional short-term sperm storage and reproduction in two oviparous sceloporine lizards from Florida, USA | https://www.researchgate.net/publication/351525317 |
| A28 | Behavioural avoidance of sperm ageing depends on genetic similarity of mates in a monogamous seabird | 10.1093/biolinnean/blz079 |
| A29 | Multiple deleterious effects of experimentally aged sperm in a monogamous bird | 10.1073/pnas.0803067105 |
| A30 | Application of ultrasound technique to evaluate the testicular function and its correlation to the sperm quality after different collection frequency in rams | 10.3389/fvets.2022.1035036 |
| B31 | Ageing of rabbit spermatozoa in the male tract and its effect on fertility | https://doi.org/10.1530/jrf.0.0200287 |
| B32 | Effect of sexual rest and frequency of ejaculation on sperm acrosomal morphology | https://doi.org/10.3168/jds.S0022-0302(71)85879-4 |
| B33 | Effects of the ageing of rabbit spermatozoa in utero on fertilization and prenatal development | https://doi.org/10.1530/jrf.0.0200299 |
| B35 | Sperm aging in the male after sexual rest: Contribution to chromosome anomalies | https://doi.org/10.1002/mrd.1120120206 |
| B36 | Sperm morphology is better in the second ejaculate than in the first in domestic cats electroejaculated twice during the same period of anesthesia | https://doi.org/10.1016/S0093-691X(97)00048-4 |
| B37 | Pre-but not post-meiotic senescence affects sperm quality and reproductive success in the North African houbara bustard | 10.3389/fevo.2022.977184 |
| B40 | Functional sperm storage duration in female *Hemidactylus frenatus* (family Gekkonidae) |  |
| B41 | Impact of immune activation on stored sperm viability in ant queens | 10.1098/rspb.2018.2248 |
| B42 | Male age alone predicts paternity success under sperm competition when effects of age and past mating effort are experimentally separated | 10.1098/rspb.2021.0979 |
| B43 | Measuring sperm viability over time in honey bee queens to determine patterns in stored-sperm and queen longevity | 10.3896/IBRA.1.53.4.02 |
| B45 | Prudent sperm use by leaf-cutter ant queens | 10.1098/rspb.2009.1184 |

**Other supplementary figures**

**A**

**B**
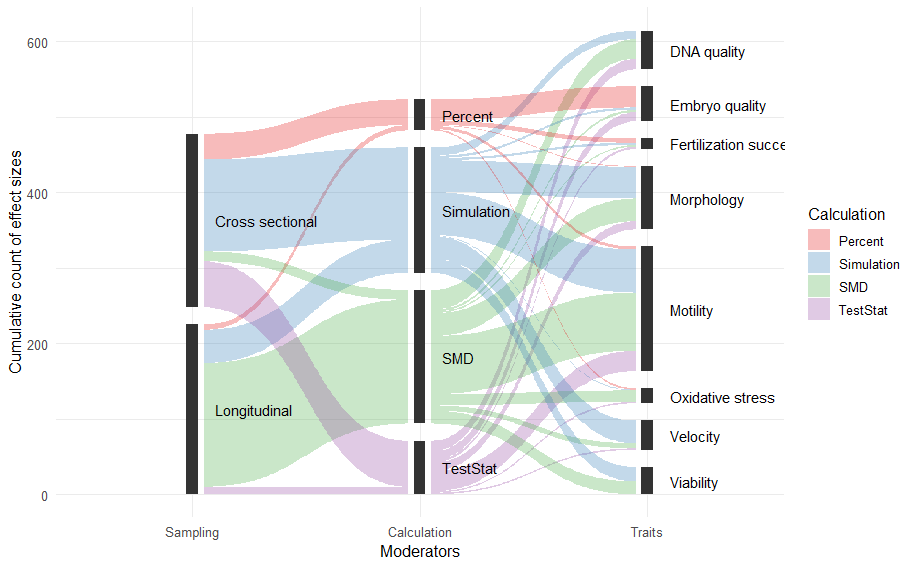

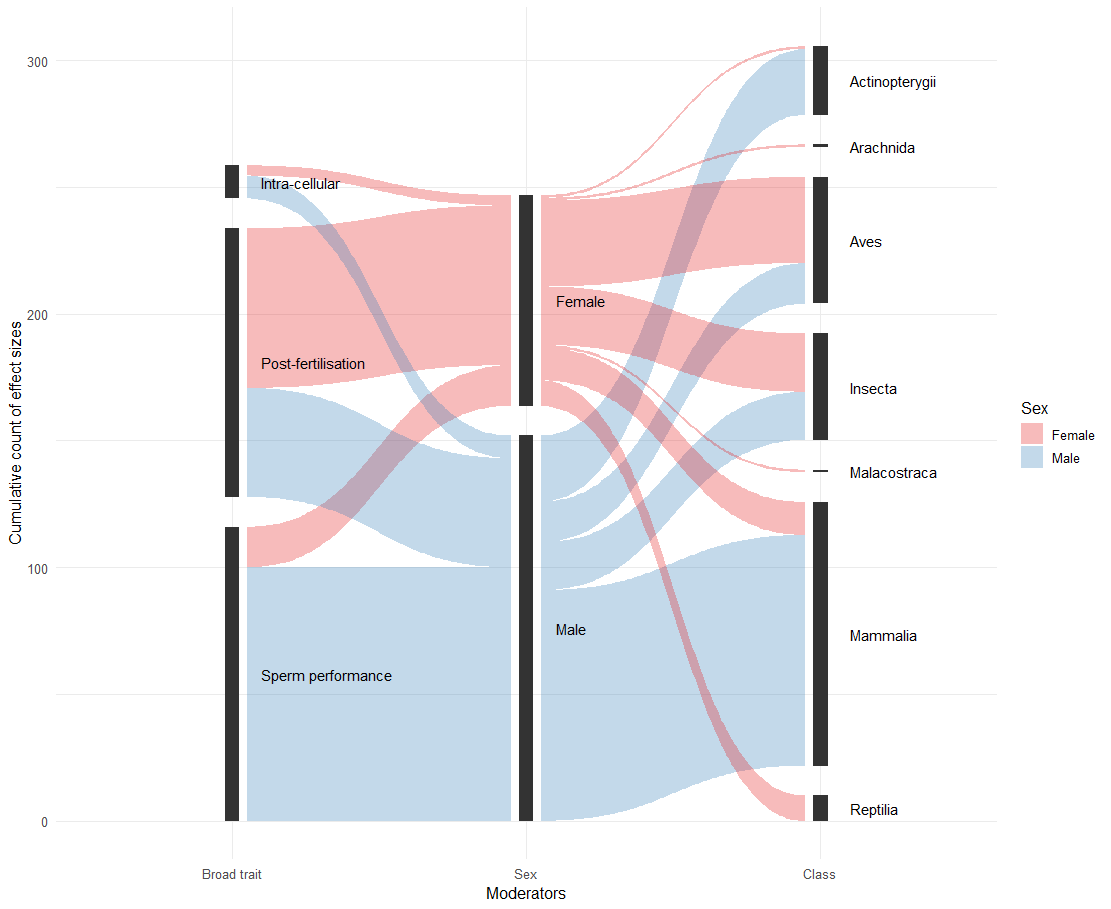

Figure S4: Alluvial plot representing the distribution of effect sizes across various moderators and each of their levels in the **(A)** human and **(B)** animal dataset. Visually, there is collinearity between effect size calculation method and male sampling, with most effect sizes from longitudinal studies being calculated as SMD. There is collinearity between trait type and sex of sperm storage, with most data on sperm performance traits being obtained from studies with male storage.

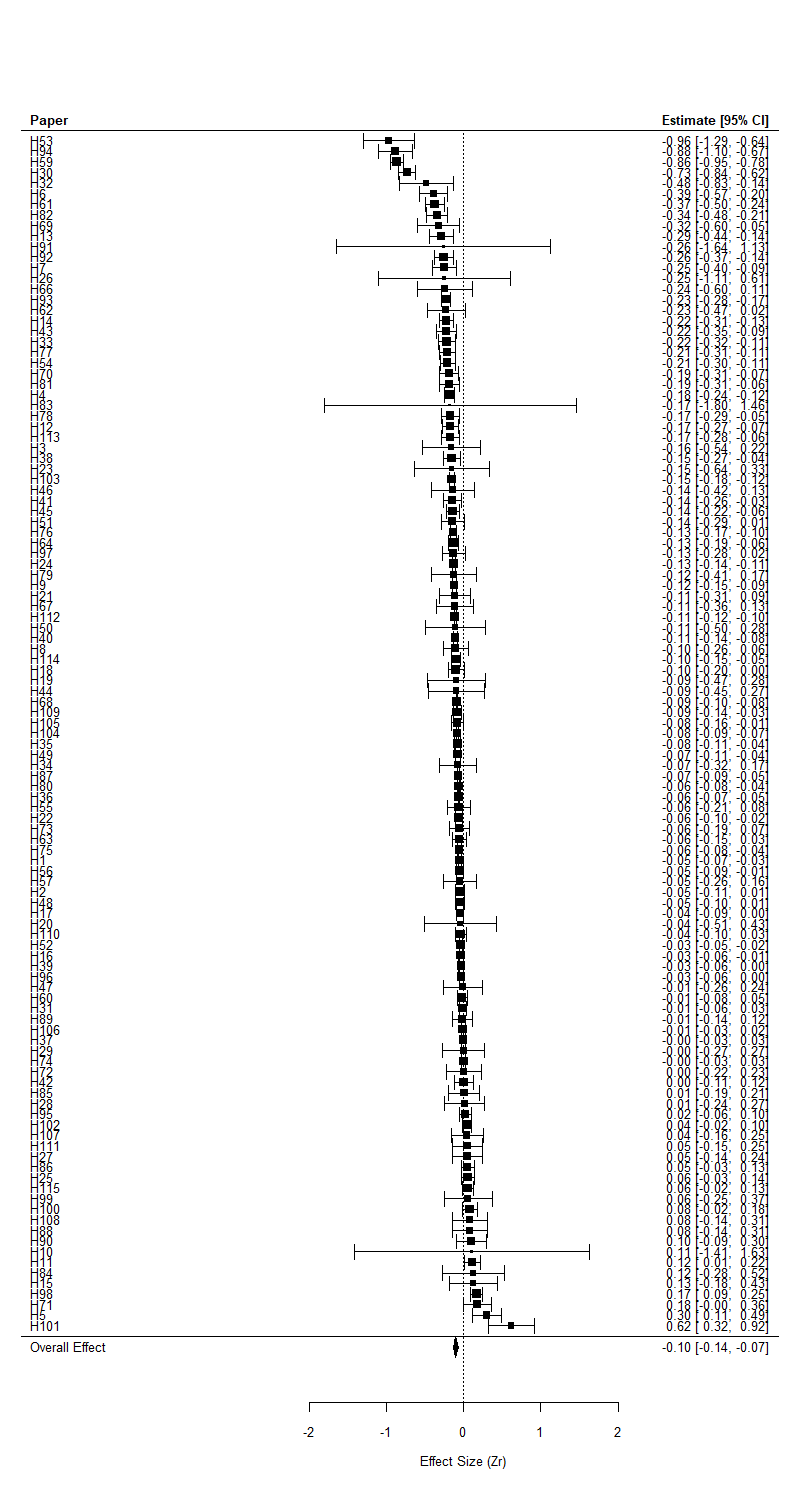

Figure S5: Mean effect size (Zr) per study for human dataset. Dark square shows mean of the study weighted-averaged across multiple effect sizes, error bars show 95% C.I. “Overall Effect” diamond shows the unweighted mean of effect sizes across all studies.

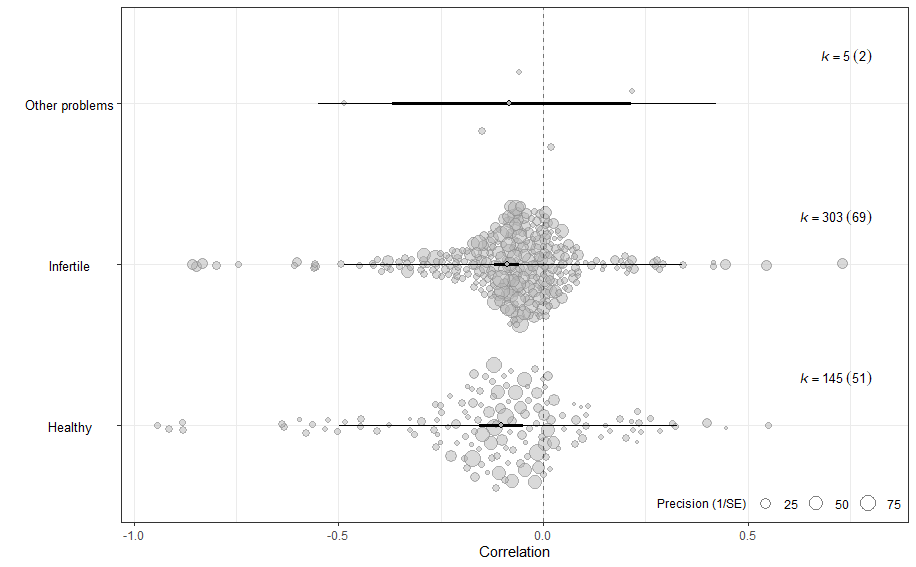

Figure S6: Health condition of males did not significantly modulate outcomes in humans. Negative correlations (Zr back-transformed to Pearson’s r) depict poorer reproductive outcomes when sperm are stored, and positive values represent improvements in reproductive outcomes with storage. The size of each datapoint represents precision of the effect size. Bold error bars show confidence intervals, and light bars show prediction intervals. Samples sizes k = number of effect sizes (number of studies in brackets).

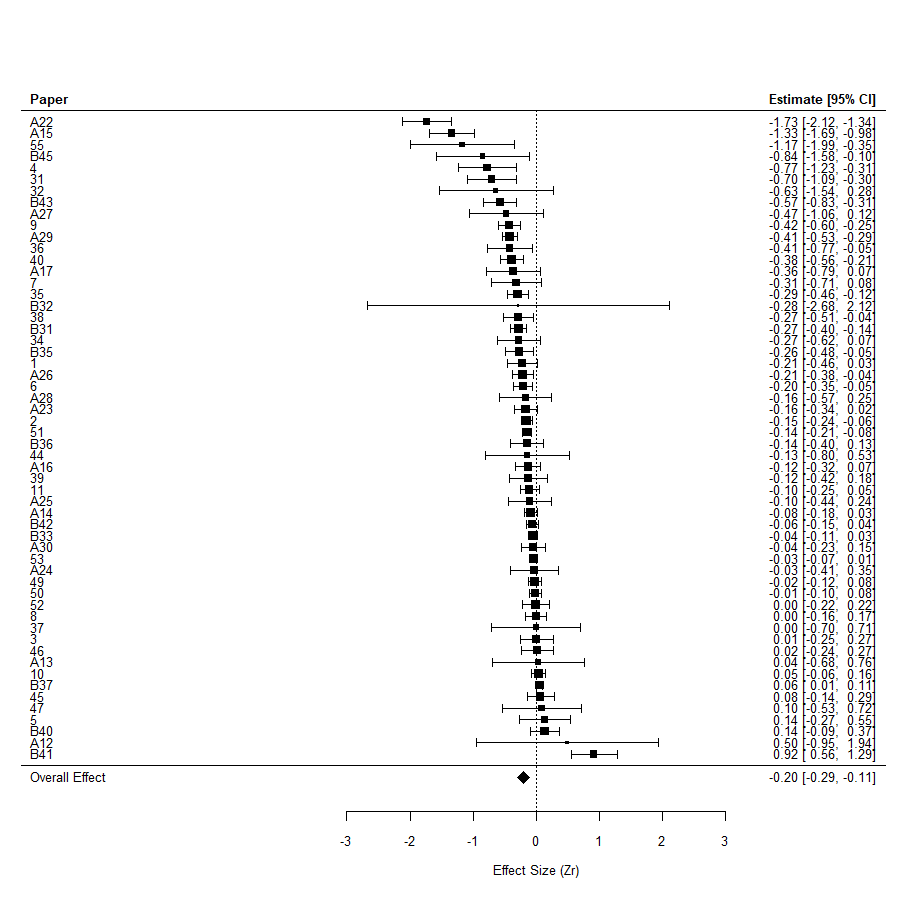

Figure S7: Mean effect size (Zr) per study for animal dataset. Dark square shows mean of the study weighted-averaged across multiple effect sizes, error bars show 95% C.I. “Overall Effect” diamond shows the unweighted mean of effect sizes across all studies.

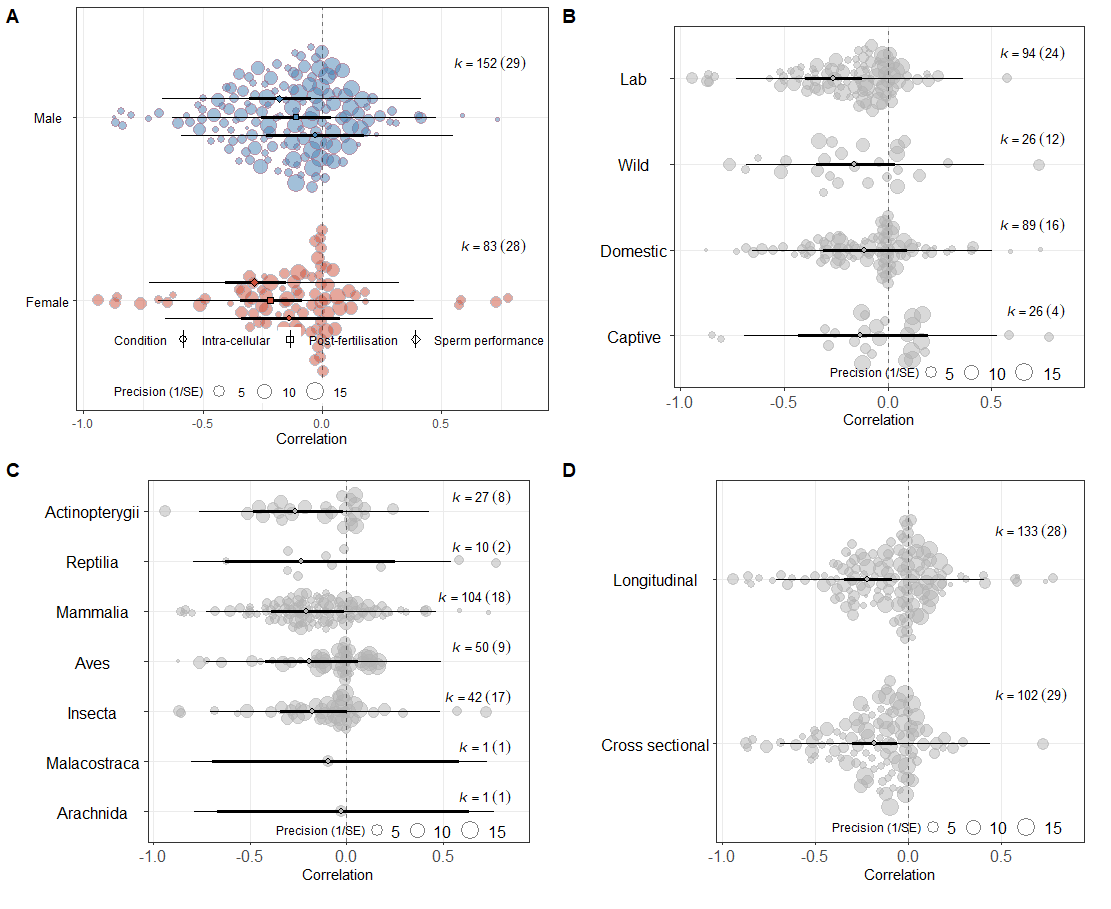

Figure S8: Influence of **(A)** trait-type * sex, **(B)** population setting, **(C)** taxonomic class, **(D)** sampling method of individuals, on modulating outcomes of sperm storage. Negative correlations (Zr back-transformed to Pearson’s r) depict poorer reproductive outcomes when sperm are stored, and positive values represent improvements in reproductive outcomes with storage. The size of each datapoint represents precision of the effect size. Bold error bars show confidence intervals, and light bars show prediction intervals. Samples sizes k = number of effect sizes (number of studies in brackets).

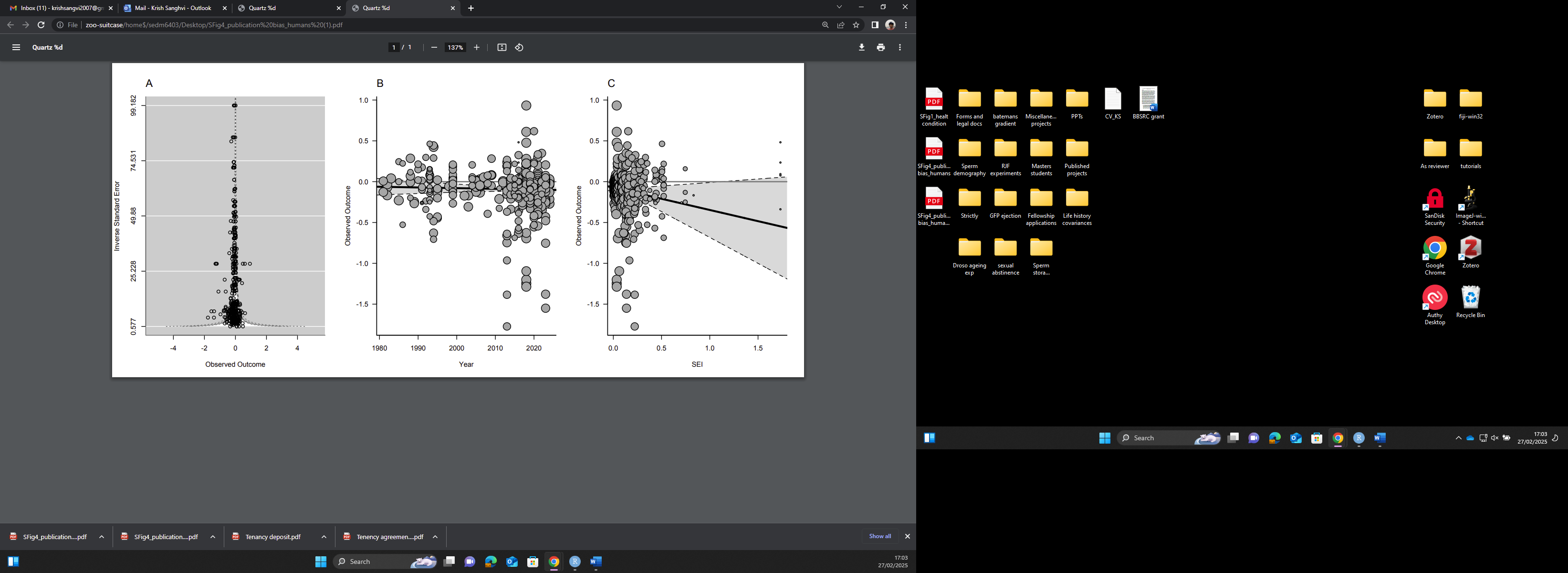

Figure S9: No evidence for publication bias in the human dataset, assessed by inspecting **(A)** funnel plots, **(B)** via time-lag bias, and **(C)** small study bias analyses. In B and C, size of points represents precision, shaded areas represent 95% C.I. Negative observed outcome values (Zr) depict poorer reproductive outcomes when sperm are stored, and positive values represent improvements in reproductive outcomes with storage.

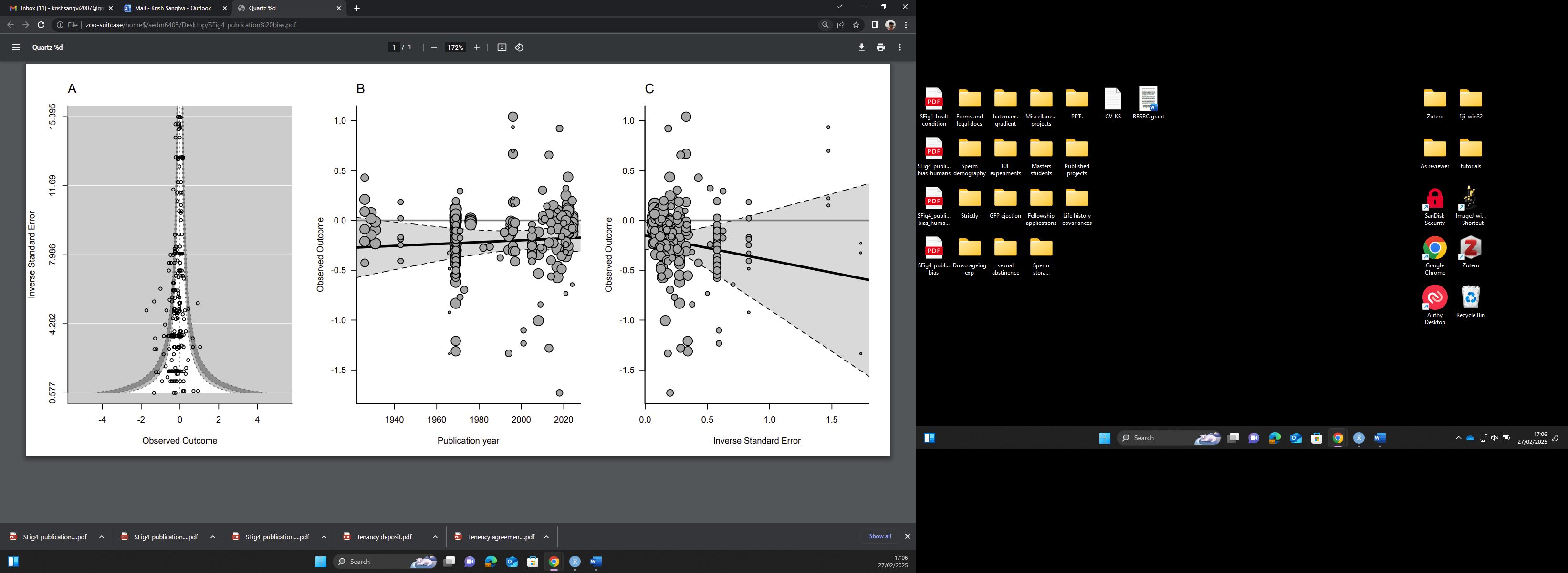

Figure S10: No evidence for publication bias in the animal dataset, assessed by inspecting **(A)** funnel plots, **(B)** via time-lag bias, and **(C)** small study bias analyses. In B and C, size of points represents precision, shaded areas represent 95% C.I. Negative observed outcome values (Zr) depict poorer reproductive outcomes when sperm are stored, and positive values represent improvements in reproductive outcomes with storage.
